# Tri-Modal Single-Cell Atlas of Developing Human Brain

**DOI:** 10.64898/2026.11.10.765750

**Authors:** Liting Song, Zhe Wang, Bohan Zhu, Faith A. DeKuyper, Cheng Wang, Zhicong Liao, Kiran Girdhar, Donghoon Lee, Nadejda M. Tsankova, Deepak A. Kaji, John F. Fullard, Pengfei Dong, Panos Roussos, Sai Ma

**Affiliations:** Center for Disease Neurogenomics, Icahn School of Medicine at Mount Sinai, New York, NY 10029, USA; Friedman Brain Institute, Icahn School of Medicine at Mount Sinai, New York, NY 10029, USA; Department of Psychiatry, Icahn School of Medicine at Mount Sinai, New York, NY 10029, USA; Department of Genetics and Genomic Sciences, Icahn School of Medicine at Mount Sinai, New York, NY 10029, USA; Department of Pathology, Molecular, and Cell-Based Medicine, Icahn School of Medicine at Mount Sinai, New York, NY 10029, USA; Department of Neuroscience, Icahn School of Medicine at Mount Sinai, New York, NY 10029, USA; Center for Precision Medicine and Translational Therapeutics, James J. Peters VA Medical Center, Bronx, NY 10468, USA; Mental Illness Research, Education and Clinical Center VISN2, James J. Peters VA Medical Center, Bronx, NY 10468, USA; Institute of Genomic Health, Icahn School of Medicine at Mount Sinai, New York, NY 10029, USA; Department of Radiation Oncology, Shandong Cancer Hospital and Institute, Shandong First Medical University and Shandong Academy of Medical Sciences, Jinan, Shandong Province 250117, China

**Keywords:** Human brain development, Single-cell multi-omics, DNA methylation dynamics, Regulatory network

## Abstract

Epigenetic modifications, such as DNA methylation, evolve throughout development and aging, providing remarkable regulatory diversity and contributing to disease etiology, including neuropsychiatric disorders. However, how these regulatory layers interact across human brain development remains unclear. Leveraging ME-seq, we simultaneously mapped gene expression, chromatin accessibility, and DNA methylation in over 340,000 single nuclei from 40 human brain specimens spanning fetal development to adolescence. Chromatin accessibility and DNA methylation captured partially overlapping *cis*-regulatory associations. Developmental trajectories revealed distinct temporal kinetics: DNA methylation continued remodeling beyond major transcriptional and accessibility transitions, while developmental clocks yielded modality-dependent age estimates. Modality-specific regulatory networks converged on developmental transcription factors, whereas methylation-defined regulatory regions provided complementary annotations of neuropsychiatric genetic risk. Together, these findings establish DNA methylation as a persistent and complementary dimension of developmental regulation, linking prolonged remodeling to human brain maturation and disease susceptibility.

**Highlights:**

- A single-cell tri-modal atlas across human brain development
- DNA methylation and accessibility define complementary *cis*-regulatory architectures
- DNA methylation remodeling persists beyond transcriptional and accessibility changes
- Developmental methylation and accessibility capture distinct neuropsychiatric risk loci

**Graphical abstract:** 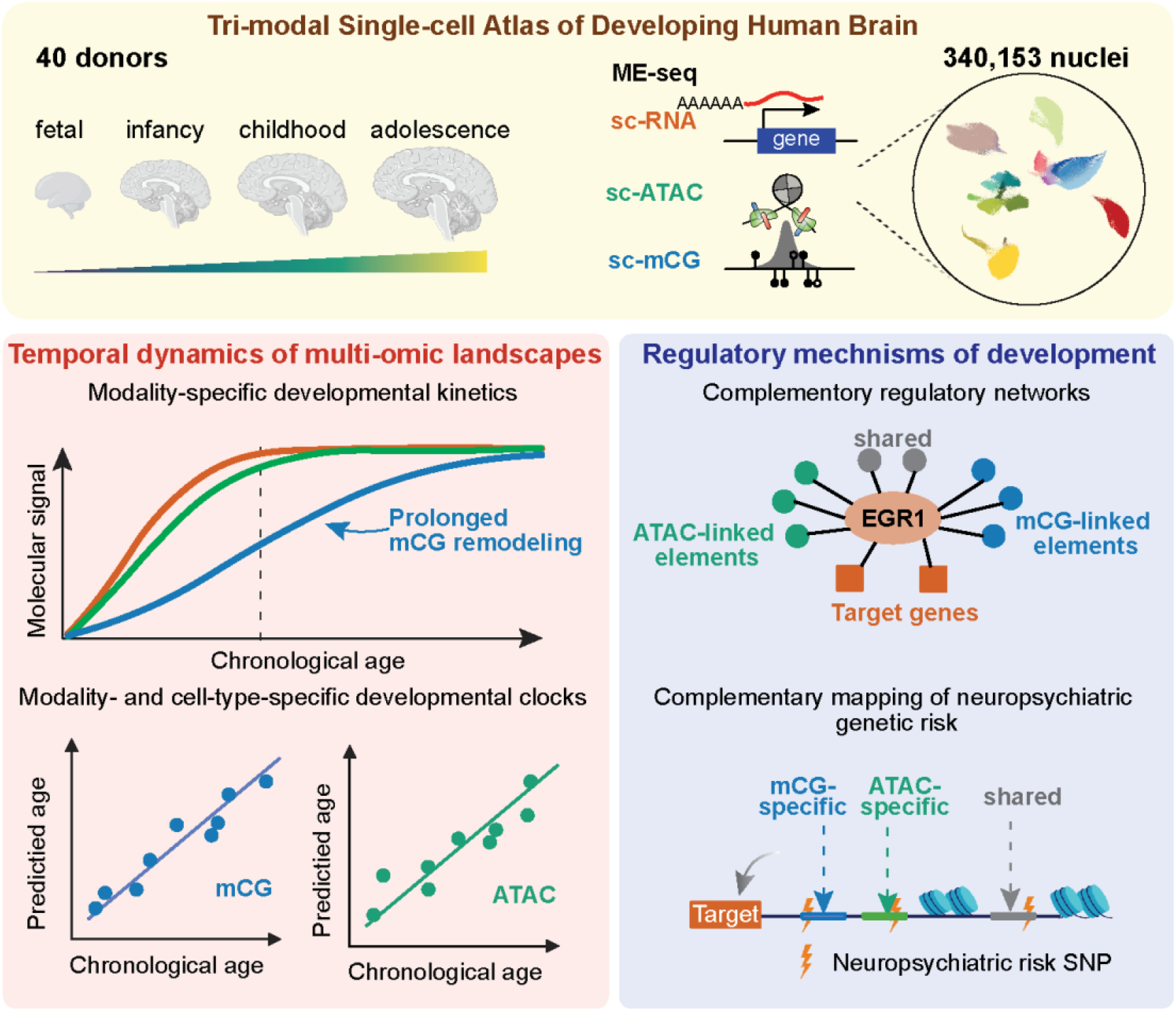

## Introduction

Human brain development unfolds over an exceptionally prolonged period beyond prenatal neurogenesis through childhood and adolescence, during which progenitor cells give rise to diverse neuronal and glial populations that progressively acquire specialized molecular and functional identities ^1–3^. Establishing these identities requires extensive and coordinated remodeling of transcription and underlying epigenetic landscapes, involving DNA methylation and chromatin accessibility ^4–7^. Yet cellular maturation is not a single, synchronized molecular process. Transcriptional programs can change rapidly during cell fate transitions, whereas epigenetic states may evolve over longer developmental intervals ^8,9^. Do these molecular layers follow a common developmental trajectory, or do they encode distinct regulatory information and operate on different timescales? Resolving this question is essential for understanding the molecular principles governing human brain maturation.

Recent single-cell atlases have transformed our understanding of molecular dynamics across human brain development, revealing pronounced cell-type-specific regulatory relationships between transcription, chromatin accessibility, and DNA methylation ^8,10–13^. However, concordance in cellular identity does not necessarily imply equivalence in regulatory information. Chromatin accessibility identifies genomic regions permissive to regulatory factor binding, whereas DNA methylation reflects an additional dimension of epigenetic regulation that can vary independently of chromatin accessibility ^9,14^. Whether these modalities delineate the same *cis*-regulatory relationships or capture complementary regulatory architectures across human brain development remains incompletely understood.

These unresolved relationships are also critical for dissecting the genetic basis of neuropsychiatric diseases. Disease-associated variants are enriched in non-coding regulatory regions associated with specific brain cell types and developmental periods ^15^, yet most functional annotations rely heavily on chromatin accessibility or transcriptional measurements. Should DNA methylation capture regulatory states that are not fully represented by accessible chromatin, developmental methylation maps could reveal additional disease-relevant regulatory elements and connect genetic risk to molecular programs that would otherwise remain unresolved.

Resolving these questions requires direct comparison of transcriptional and epigenetic regulation within the same cells. However, most existing developmental atlases rely on independently generated datasets, where differences in cellular composition, developmental sampling, and assay sensitivity complicate cross-modal comparisons ^16,17^. Simultaneous profiling of gene expression, chromatin accessibility, and DNA methylation provides a unified framework for dissecting their regulatory relationships and temporal dynamics throughout human brain development.

Here, we leveraged ME-seq (**M**ethylation and **E**xpression co-assay with **seq**uencing) ^18^ to generate a single-cell tri-modal atlas of the developing human brain, jointly profiling gene expression, chromatin accessibility, and CpG methylation (mCG) from fetal life through adolescence. Across 340,153 nuclei from 40 donors, we reconstructed major neuronal and glial developmental trajectories and directly compared the shared and modality-specific *cis*-regulatory relationships across the three regulatory layers. We find that chromatin accessibility and mCG define partially overlapping sets of regulatory architectures. Moreover, although the three modalities are broadly coordinated during differentiation, they exhibit distinct developmental kinetics: mCG undergoes prolonged remodeling across postnatal excitatory neuronal and oligodendroglial maturation, extending beyond major transcriptional and accessibility transitions, with developmental dynamics further supported by cortical organoid models and independent human brain datasets. Integrative regulatory-network analyses identify shared developmental transcription factors (TFs) acting through partially distinct epigenomic architectures, while genetic analyses show that methylation-defined regulatory elements capture neuropsychiatric genetic risk not fully resolved by chromatin accessibility. Together, these findings reveal that human brain maturation is organized across regulatory layers with distinct temporal and functional properties and provide a framework for connecting developmental epigenomic remodeling to disease susceptibility.

## Results

### A tri-modal single-cell atlas of human brain development

To define transcriptomic and epigenomic remodeling across human brain development, we collected 40 brain specimens spanning fetal development, infancy, childhood, and adolescence (**Fig. 1A, Table S1**). Using ME-seq, we generated 340,153 high-quality single-nucleus profiles jointly capturing the transcriptome, chromatin accessibility, and mCG after quality filtering (**Fig. S1A**), creating a tri-modal single-cell atlas of the developing human brain.

**Figure 1.**
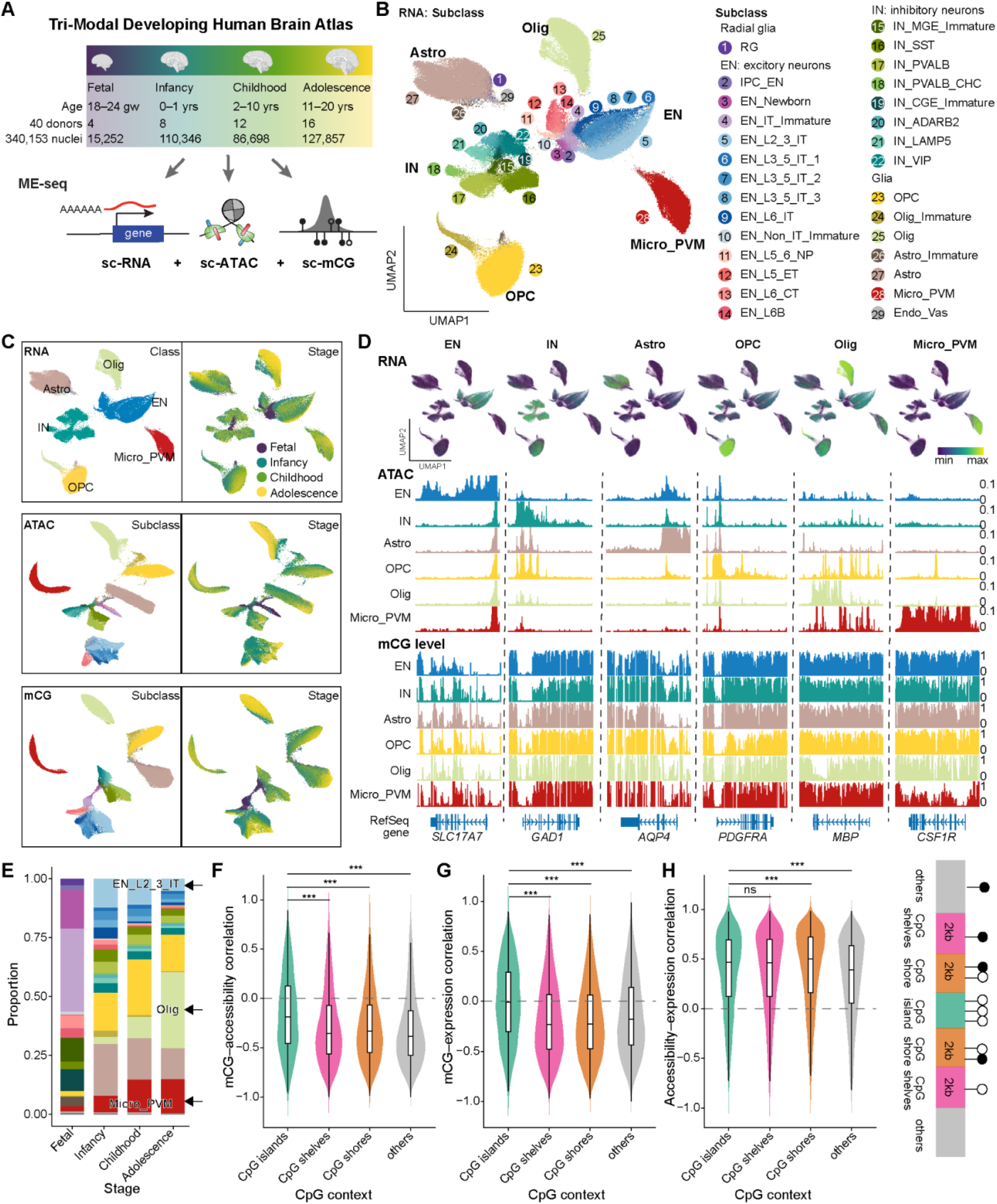
A tri-modal single-cell atlas of human brain development. **(A)** Overview of the study design. Human brain samples from 40 donors spanning fetal development through adolescence were profiled using ME-seq to jointly measure gene expression, chromatin accessibility, and DNA methylation in 340,153 single nuclei. Numbers indicate the number of donors and nuclei at each developmental stage. gw, gestational week; yrs, years. **(B)** UMAP of the RNA profiles from ME-seq showing 29 annotated cell subclasses, including radial glia (RG) and intermediate progenitor cells (IPCs), newborn and immature excitatory neurons (ENs), medial ganglionic eminence (MGE)- and caudal ganglionic eminence (CGE)-derived inhibitory interneurons (INs), microglia and perivascular macrophages (Micro_PVMs), oligodendrocyte precursor cells (OPCs), immature and mature oligodendrocytes (Olig), astrocytes (Astro), and endothelial and vascular cells (Endo_vas). **(C)** UMAP embeddings of RNA (top), ATAC (middle), and mCG (bottom) profiles. RNA profiles are colored by major cell class (left) and developmental stage (right); ATAC and mCG profiles are colored by cell subclass (left) and developmental stage (right). **(D)** Representative canonical cell-type marker genes across three modalities. **(E)** Relative cellular composition across fetal, infancy, childhood, and adolescent stages. **(F–H)** Violin plots showing distributions of pairwise correlations among DNA methylation, chromatin accessibility, and gene expression at cell-type-specific DMRs, stratified by genomic CpG context. Boxplots indicate medians and interquartile ranges. Correlations were calculated using Spearman’s rank correlation coefficient. Differences between CpG contexts were assessed using the Wilcoxon test. \*\*\**FDR* < 0.001; ns, not significant.

Using the RNA profiles of ME-seq data, we annotated 29 subclasses across eight major cell classes (**Fig. 1B, Fig. S1B**), encompassing progenitors, immature and mature neuronal populations, and major glial and vascular cell types. Independent embeddings based on chromatin accessibility and mCG resolved highly concordant cellular populations (**Fig. 1C, Fig. S1C**), supporting the reproducibility of cell-type identification across modalities. Consistent with established gene regulatory relationships ^19^, canonical marker genes exhibited concordant molecular signatures across modalities, with cell-type-specific expression accompanied by increased chromatin accessibility and reduced mCG levels in their corresponding cell types (**Fig. 1D, Fig. S1B**). Across all three modalities, cell embeddings were organized primarily by cell type and developmental stage rather than technical batch, indicating effective control of batch effects. Cellular composition shifted substantially across development, with progenitors and immature neurons declining and glial populations, particularly microglia and oligodendrocytes, increasing postnatally (**Fig. 1E**).

We next characterized the relationships among the three molecular modalities across genomic contexts at cell-type-specific differentially methylated regions (DMRs; **Table S2**). Chromatin accessibility and gene expression showed broadly positive correlations, whereas methylation-associated correlations tended to be negative, with their strength varying by local CpG context (**Fig. 1F–H**). Specifically, mCG showed relatively weak correlations with both accessibility and gene expression at CpG islands, but more pronounced inverse correlations in CpG shores, shelves, and other non-island regions (**Fig. 1F, G**). These patterns are consistent with the generally low methylation levels of promoter-associated CGIs and suggest that methylation variation outside CpG islands may more closely reflect regulatory activity. Together, these findings highlight the context-dependent relationships among the three modalities and motivate a direct comparison of the *cis*-regulatory associations captured by chromatin accessibility and DNA methylation.

### Chromatin accessibility and DNA methylation define complementary *cis*-regulatory architectures

Both DNA methylation and chromatin accessibility shape gene expression during cortical development, but whether they converge on the same *cis*-regulatory relationships is unclear. Despite advances in bimodal ATAC–RNA or mCG–RNA mapping ^20,21^, their relative regulatory contributions remain incompletely defined. Leveraging the matched tri-modal measurements, we directly compared ATAC- and mCG-derived *cis*-relationships within a common genomic framework. We partitioned ±250 kb genomic windows around transcription start sites (TSSs) into 500-bp tiles and identified 83,698 ATAC–RNA and 71,847 mCG–RNA putative regulatory links within ±50 kb of TSSs across the full dataset and cell-type-stratified analyses, leveraging the covariations between gene expression and accessibility or methylation levels across single cells (**Fig. 2A**, **Fig. S2A–D**, **Table S3**, **S4**; see STAR Methods). These regulatory links further identified 801 ATAC-derived and 1,147 mCG-derived domains of regulatory chromatin (DORC) genes ^20^ with a large number of associated regulatory elements (**Fig. 2B**, **C**). Top-ranked DORC genes were largely shared between the two modalities, including canonical cortical regulators such as *SATB2*, *BCL11A*, *MEF2C*, *DLX5*, and *NEUROD6*.

**Figure 2.**
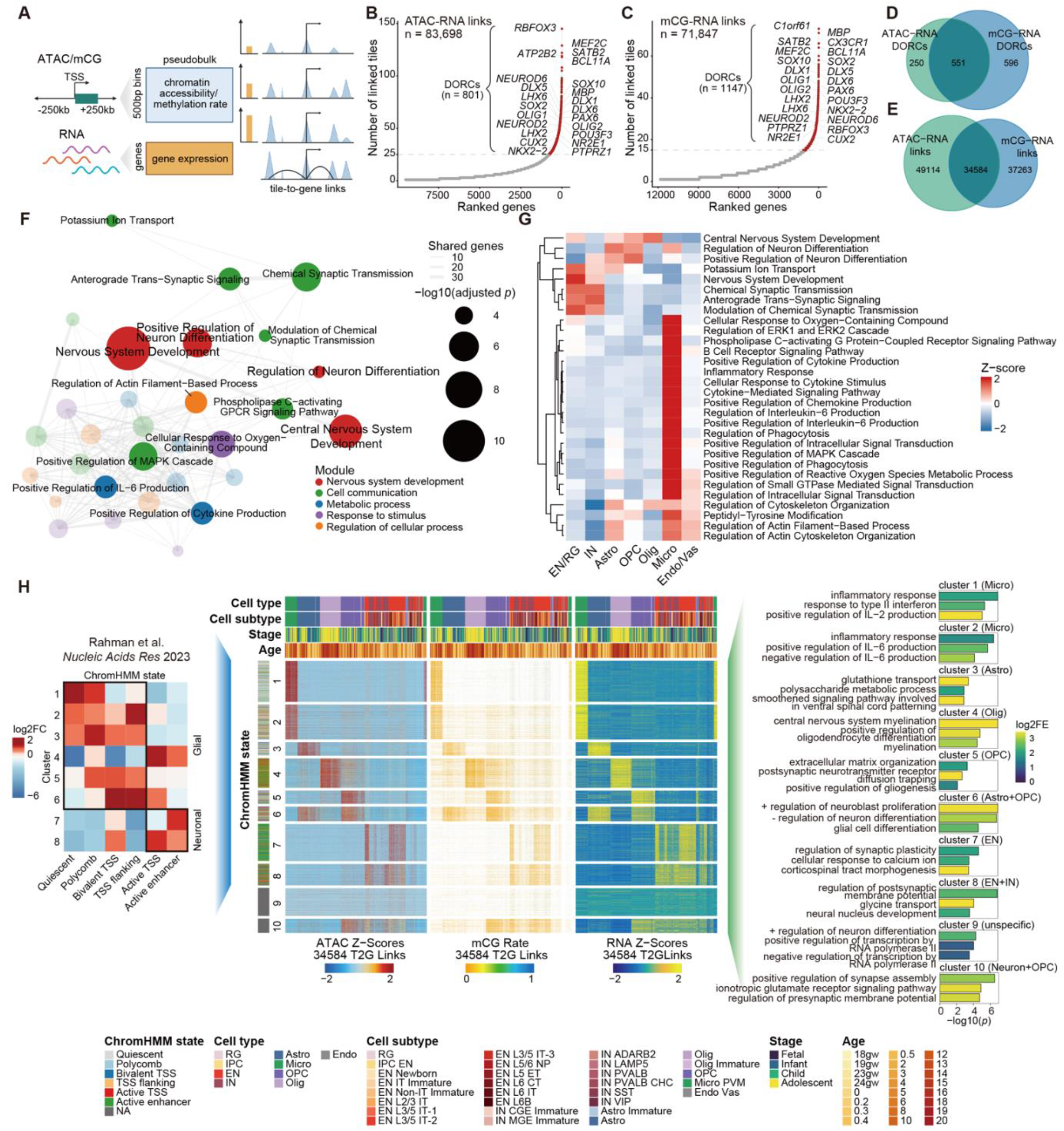
Chromatin accessibility and DNA methylation define complementary *cis*-regulatory architectures. **(A)** Schematic of the tile-based *cis*-regulatory analysis framework. Protein-coding gene TSSs were flanked by ±250 kb and partitioned into 500-bp tiles for initial association mapping, with significant links subsequently restricted to ±50 kb for DORC analyses. Tile–gene Spearman correlations were computed across matched pseudobulks and tested against a covariate-matched background for ATAC–RNA and mCG–RNA. DORCs were defined based on the number of significant tile–gene links within ±50 kb of the TSS. **(B)** Ranked distribution of significant tile–gene link numbers per gene. Significant links from the full dataset and cell-type-stratified analyses were pooled and deduplicated. Genes with more than 25 significantly associated tiles were designated as ATAC–RNA DORCs (n = 801). Top genes are labeled. **(C)** As in (B), for mCG–RNA associations. Genes with ≥ 15 significantly linked tiles were designated as mCG–RNA DORCs (n = 1,147). **(D, E)** Venn diagrams showing modality-specific and shared DORC genes (D) and significant tile–gene links (E) between ATAC–RNA and mCG–RNA. **(F)** Gene Ontology (GO) biological process enrichment network of the unified DORC gene set derived from both ATAC and mCG. Each node represents one of the top 30 terms ranked by adjusted *p*-value, with size indicating −log_10_(adjusted *p*-value) and color indicating functional modules. Edges connect terms with shared genes, with width proportional to shared gene counts. **(G)** Heatmap of module scores for the top 30 enriched GO terms across cell types. Z-scores are standardized across cell types within each term. Rows are hierarchically clustered. **(H)** Middle: Heatmap of chromatin accessibility, DNA methylation, and gene expression for 34,584 jointly recovered tile–gene links grouped into 10 k-means clusters. Left: ChromHMM state enrichment using glial segmentations for clusters 1–6 and neuronal segmentations for clusters 7–8. Clusters 9–10 were excluded. Right: top GO biological process terms enriched among the 200 most frequently linked genes per cluster. log2FE, log2 fold enrichment.

Despite convergence on these core regulators, the two epigenetic modalities recovered substantial non-overlapping regulatory information: 551 DORCs were shared with moderate concordance in gene rankings (Spearman’s ρ = 0.486, *p* < 2.2e-16), whereas 250 were specific to ATAC and 596 to mCG; similarly, only 34,584 individual putative regulatory links were recovered by both modalities (**Fig. 2D**, **E**). Notably, the modality-specific DORCs were enriched for distinct functional categories. mCG-specific DORCs included GABAergic, glial, and innate immune functions, whereas ATAC-specific DORCs were enriched for synaptic transmission and ion channel processes (**Table S5**, **S6**). Importantly, the linked tiles exhibited elevated evolutionary conservation (**Fig. S2E**) and DORC genes were strongly enriched for super-enhancer-associated genes ^15,22^ (**Fig. S2F**, **G**), supporting the regulatory relevance of these associations.

Functional annotation of the unified DORC gene set derived from accessibility and methylation revealed distinct regulatory modules associated with nervous system development, cell communication, metabolism, stimulus responses, and regulation of cellular processes (**Fig. 2F**). These modules exhibited distinct activity patterns across cell types (**Fig. 2G**), highlighting the organization of *cis*-regulatory programs across the cellular diversity of the developing cortex. Across the 34,584 shared links, increased accessibility was generally accompanied by reduced DNA methylation and increased expression of linked genes (**Fig. 2H**). However, the underlying regulatory architecture was organized into distinct cell-type-specific programs. Neuronal links were preferentially associated with active TSS and enhancer states, whereas glial links encompassed a broader spectrum of active, poised, and Polycomb-associated chromatin states. Corresponding target genes were enriched for lineage-appropriate functions, including synaptic processes in excitatory neurons, myelination in oligodendroglial cells, and inflammatory responses in microglia (**Fig. 2H**). Within ENs (**Fig. S2H**, **I**), regulatory associations further separated fetal and postnatal programs. Fetal-enriched links were preferentially associated with TSS-proximal chromatin states, whereas postnatal links showed greater enrichment for active enhancers (**Fig. S2J**), consistent with a shift toward enhancer-associated regulation during neuron maturation.

Together, these findings demonstrate that chromatin accessibility and DNA methylation converge on core developmental regulators while capturing distinct and cell-type-specific *cis*-regulatory relationships. This raises the question of whether these complementary regulatory layers also exhibit distinct temporal dynamics during human brain maturation.

### DNA methylation remodeling persists beyond transcriptional and accessibility transitions

To investigate the temporal coordination of transcriptional and epigenetic remodeling during EN development, we independently reconstructed developmental trajectories from gene expression, chromatin accessibility, and mCG profiles. All three modalities consistently captured progression from radial glia (RG) through intermediate progenitor cells (IPCs), newborn and immature ENs, to layer-specific mature ENs (**Fig. 3A, Table S7**). Across all three modalities, pseudotime increased with chronological age, with lineage- and modality-specific patterns (**Fig. 3B, Fig. S3A, B**). Specifically, deep-layer ENs showed limited progression beyond infancy, whereas upper-layer L2/3 IT neurons continued to progress through childhood and adolescence (**Fig. 3B, Fig. S3A, B**). Notably, mCG- and ATAC-derived trajectories showed a greater fraction of pseudotime progression after infancy than RNA-derived trajectories (**Fig. 3C, D**). These findings suggest lineage- and modality-specific developmental kinetics during EN maturation.

**Figure 3.**
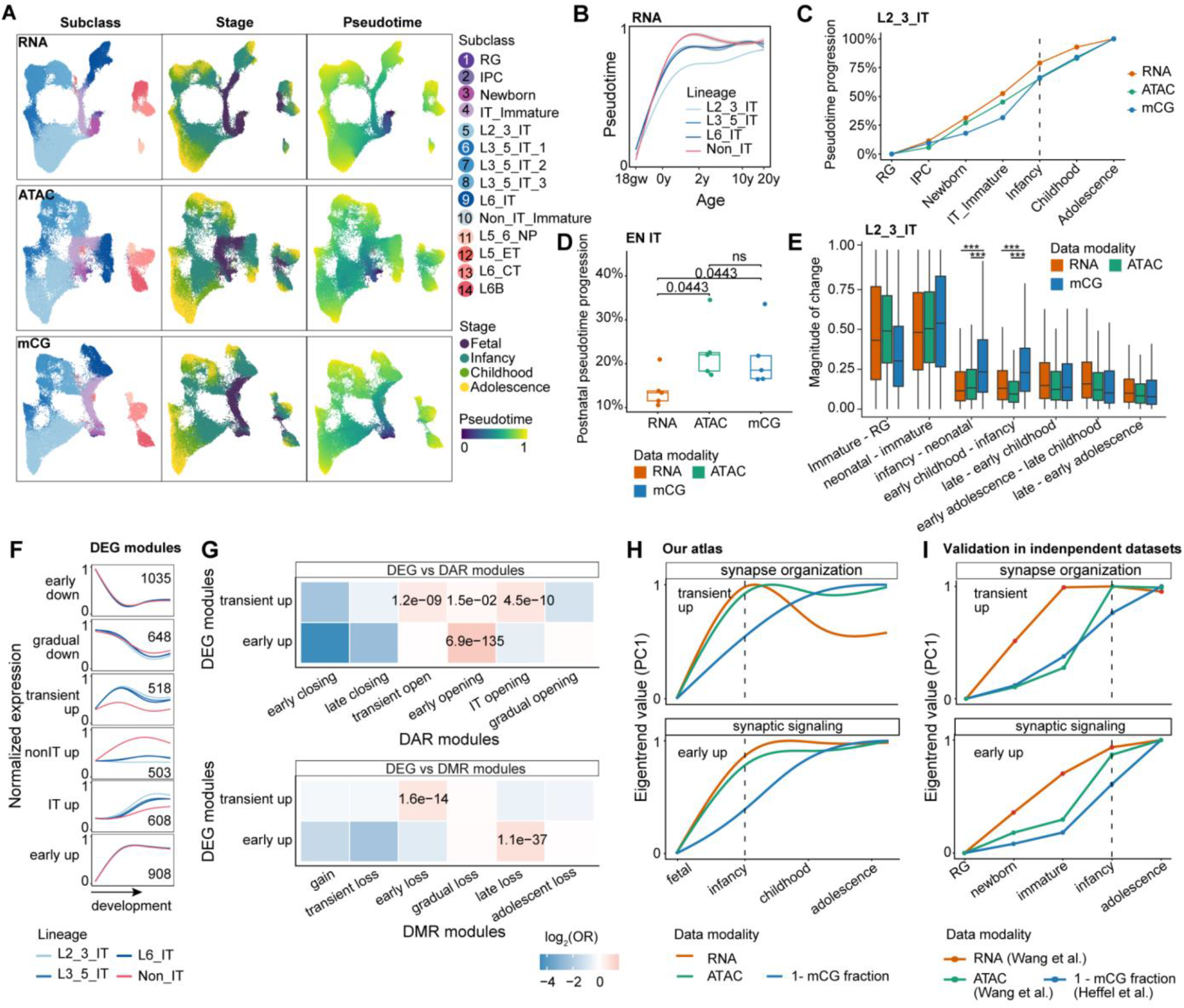
Temporal separation between transcriptional and epigenetic remodeling during EN maturation. **(A)** Trajectory inference of EN development in the RNA (top), ATAC (middle), and mCG (bottom) modalities. UMAPs are colored by subclass (left), developmental stage (middle), and inferred pseudotime (right). **(B)** Relationships between RNA-derived pseudotime and age across EN lineages. Curves were fitted using generalized additive models (GAMs). **(C)** Cumulative pseudotime progression along the L2/3 IT neuron developmental trajectory across three modalities, normalized to the total pseudotime difference between RG and adolescent neurons. The dashed vertical line indicates infancy. **(D)** Fraction of total pseudotime progression occurring during postnatal development (infancy to adolescence) across EN intratelencephalic (IT) trajectories for each modality. Each point represents an individual trajectory, and boxplots show the median and interquartile range. Pairwise comparisons were performed using one-sided Wilcoxon signed-rank tests. *FDR* values are shown. ns, not significant. **(E)** Relative magnitude of molecular changes between consecutive developmental stages in L2/3 IT neurons, calculated for dynamic DEGs, DARs, and DMRs. Boxplots show the distributions of changes across individual features. Developmental bins included RG and IPCs, newborn and immature ENs, neonatal (birth), early infancy (0.2–0.5 years), early childhood (2–4 years), late childhood (5–10 years), early adolescence (12–14 years), and late adolescence (15–20 years). *** *FDR* < 0.001. **(F)** Mean normalized expression profiles of DEGs across developmental modules during EN maturation. **(G)** Enrichment of early-upregulated and transiently upregulated DEG modules among DAR (top) and DMR (bottom) modules, assessed using one-sided Fisher’s exact tests. Colors indicate log_2_ odds ratios (OR), and values indicate *FDR*. **(H)** Eigentrend values (PC1) of gene expression, chromatin accessibility, and unmethylated CG fraction (1 – mCG fraction) across development for synapse organization and synaptic signaling in L2/3 IT neurons. **(I)** Eigentrend values of gene expression, chromatin accessibility, and unmethylated CG fraction (1 – mCG fraction) across development in L2/3 IT neurons from independent datasets ^8,13^ for synapse organization and synaptic signaling.

To characterize the dynamic molecular changes underlying these trajectories, we next identified differentially expressed genes (DEGs), differentially accessible regions (DARs), and DMRs (**Fig. S3C–E, Table S8–S10**). Although relative molecular changes were greatest during early developmental transitions, transcriptional and chromatin accessibility changes declined more rapidly after birth, whereas DNA methylation continued to undergo substantial remodeling through infancy and childhood (**Fig. 3E**). Dynamic molecular features were further grouped into modality-specific developmental modules, representing increasing, decreasing, and transient developmental patterns (**Fig. 3F, Fig. S3C–E**). These patterns were recapitulated in independent developmental datasets (**Fig. S3F–H**) ^8,13^, and by ME-seq profiling of neocortical organoids, which captured the early developmental transcriptional and epigenetic dynamics (**Fig. S4A–F**).

Despite their distinct developmental dynamics, transcriptional and epigenetic programs were linked to neuronal development and maturation, with modality-dependent differences in their functional associations and annotations (**Fig. S5A-C**). Transcriptionally downregulated programs were associated with neuronal differentiation and projection development, whereas transcriptional upregulation and chromatin opening were associated with synaptic processes, including synapse organization and signaling (**Fig. S5A**). Notably, later methylation-loss programs were enriched for dendritic spine development, intracellular transport, and neuronal signaling, linking prolonged DNA methylation remodeling to neuronal maturation-related processes. Chromatin-opening and methylation-loss regions preferentially transitioned from the quiescent chromatin state to active enhancers (**Fig. S5B**), reinforcing a shift toward active regulatory states during neuronal maturation ^23^. Developmental DMRs were more frequently localized to CpG shores and shelves than DARs (**Fig. S5C**), indicating differences in the genomic contexts of chromatin accessibility and DNA methylation remodeling.

However, concordance in the direction of regulatory changes did not necessarily imply synchronization in developmental timing (**Fig. S5D**). Transiently upregulated genes, whose expression peaked around infancy (**Fig. 3F)**, were preferentially associated with transient chromatin opening and early DNA methylation loss (**Fig. 3G**). Consistent with this temporal difference, synapse organization genes, including *AGRN* ^24^ (**Fig. S5E, F**), showed an early expression peak during the neonatal period and infancy, whereas DNA methylation continued to decline into adolescence after the major accessibility increase (**Fig. 3H**). A similar temporal decoupling was observed for early-upregulated genes: their expression increased rapidly and stabilized after infancy (**Fig. 3F**), whereas associated regulatory regions preferentially exhibited early chromatin opening but late DNA methylation loss (**Fig. 3G**). Accordingly, synaptic signaling programs showed continued DNA demethylation through childhood and adolescence despite relatively stable gene expression and chromatin accessibility (**Fig. 3H**). Continued methylation change after major transcriptional transitions was also observed in independent human brain developmental datasets ^8,13^ (**Fig. 3I**).

Together, these findings reveal that transcriptional and epigenetic programs can undergo concordant directional changes while remodeling over different timescales. In particular, DNA methylation remodeling extends beyond major transcriptional and accessibility transitions during EN maturation, raising the possibility that prolonged epigenetic remodeling is a broader feature of human brain development.

### DNA methylation remodeling persists across oligodendroglial maturation

To determine whether prolonged DNA methylation remodeling extends beyond ENs, we next examined the oligodendroglial lineage, which undergoes extensive postnatal maturation ^25^. Trajectory inference across RNA, chromatin accessibility, and DNA methylation resolved age-associated progression within OPCs, as well as differentiation from OPCs through immature to mature oligodendrocytes (**Fig. 4A, Table S11**). Although pseudotime increased with chronological age across modalities, RNA-derived trajectories showed more limited progression from childhood to adolescence than ATAC- and mCG-derived trajectories (**Fig. 4B-C**), highlighting modality-specific developmental kinetics during oligodendroglial maturation.

**Figure 4.**
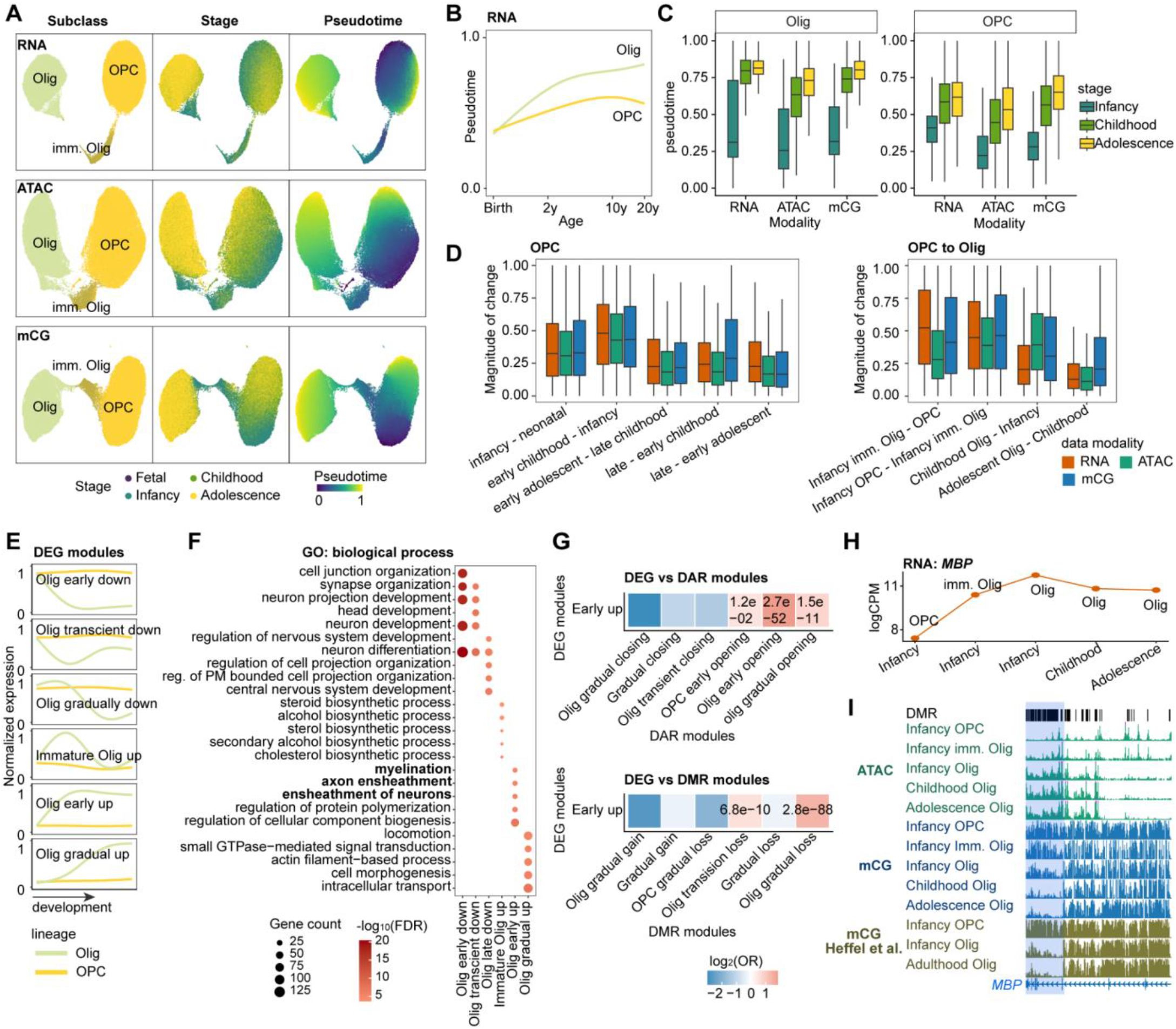
DNA methylation remodeling persists across oligodendroglial maturation. **(A)** Developmental trajectories of the oligodendroglial lineage inferred independently from RNA (top), ATAC (middle), and mCG (bottom) modalities. UMAPs are colored by subclass (left), developmental stage (middle), and inferred pseudotime (right). Imm. Olig, immature oligodendrocytes. **(B)** Relationship between RNA-derived pseudotime and age for OPC and oligodendrocytes. Curves were fitted using generalized additive models (GAMs). **(C)** Distribution of inferred pseudotime across infancy, childhood, and adolescence for oligodendrocytes and OPCs in each modality. Boxplots show the median and interquartile range. **(D)** Relative magnitude of molecular changes between consecutive developmental stages/states during OPC maturation (left) and OPC-to-oligodendrocyte differentiation (right), calculated for dynamic DEGs, DARs, and DMRs. Boxplots show the distributions of changes across individual molecular features. **(E)** Mean normalized expression profiles of DEGs in each module across development. **(F)** Gene Ontology (GO) biological process enrichment of DEG modules. (**G**) Enrichment of early-upregulated DEGs among DAR (top) or DMR (bottom) modules, assessed using one-sided Fisher’s exact tests. Colors indicate log2 odds ratios (OR), and values indicate *FDR*. **(H)** Pseudobulk expression of *MBP* (logCPM) across oligodendroglial cell types and developmental stages. **(I)** Genomic tracks of the *MBP* locus showing the chromatin accessibility and DNA methylation (mCG fraction) across oligodendroglial maturation, including mCG profiles from an independent dataset ^8^. The blue block highlights DMRs exhibiting continued DNA methylation remodeling.

Consistent with the patterns observed in ENs, transcriptional and epigenetic features were broadly coordinated but followed distinct temporal kinetics (**Fig. 4D–I, Fig. S6, Table S12–S14**). Across oligodendroglial maturation, DNA methylation continued to undergo substantial remodeling at later developmental stages, even as transcriptional and chromatin accessibility changes declined relative to early transitions (**Fig. 4D**). Although transcriptional upregulation, chromatin opening, and DNA methylation loss were all associated with oligodendroglial maturation-related processes, including myelination and axon ensheathment (**Fig. 4F, Fig. S6D, E**), their developmental changes were not temporally synchronized (**Fig. S6F**). Early-upregulated genes were preferentially associated with early chromatin opening but gradual DNA methylation loss (**Fig. 4G**). *MBP*, encoding myelin basic protein ^26^, showed markedly increased expression from infant OPCs to immature oligodendrocytes and remained elevated through adolescence (**Fig. 4H**). Chromatin accessibility increased during this transition, whereas mCG levels at the promoter-adjacent DMRs continued to decline throughout childhood and adolescence (**Fig. 4I**). Together, these findings demonstrate that DNA methylation remodeling persists beyond major transcriptional and chromatin accessibility transitions during oligodendroglial maturation, suggesting that distinct transcriptional and epigenetic timescales represent a broader feature of human brain development.

### Modality-specific developmental clocks reveal distinct temporal representations of brain maturation

The prolonged DNA methylation remodeling observed across neuronal and oligodendroglial lineages prompted us to ask whether transcriptional and epigenetic modalities encode developmental age differently. We first built modality-specific developmental clocks using LASSO regression on donor- and cell-type-stratified profiles (**Fig. 5A, Fig. S7A, B**; see STAR Methods). All three modalities accurately predicted donor age, with RNA achieving the highest accuracy (R^2^ = 0.996), followed by mCG (R^2^ = 0.959) and ATAC (R^2^ = 0.940) (**Fig. 5B**). External validations ^13,27,28^ further supported the performance of RNA- and ATAC-based clocks (**Fig. S7C**). Although the models shared a core set of genes, including *VEGFA*, *FOXO1*, *FGFR1*, and *CLU*, most selected features were modality-specific (**Fig. 5C, Fig. S7D**), suggesting that developmental timing was encoded through overlapping but distinct regulatory programs.

**Figure 5.**
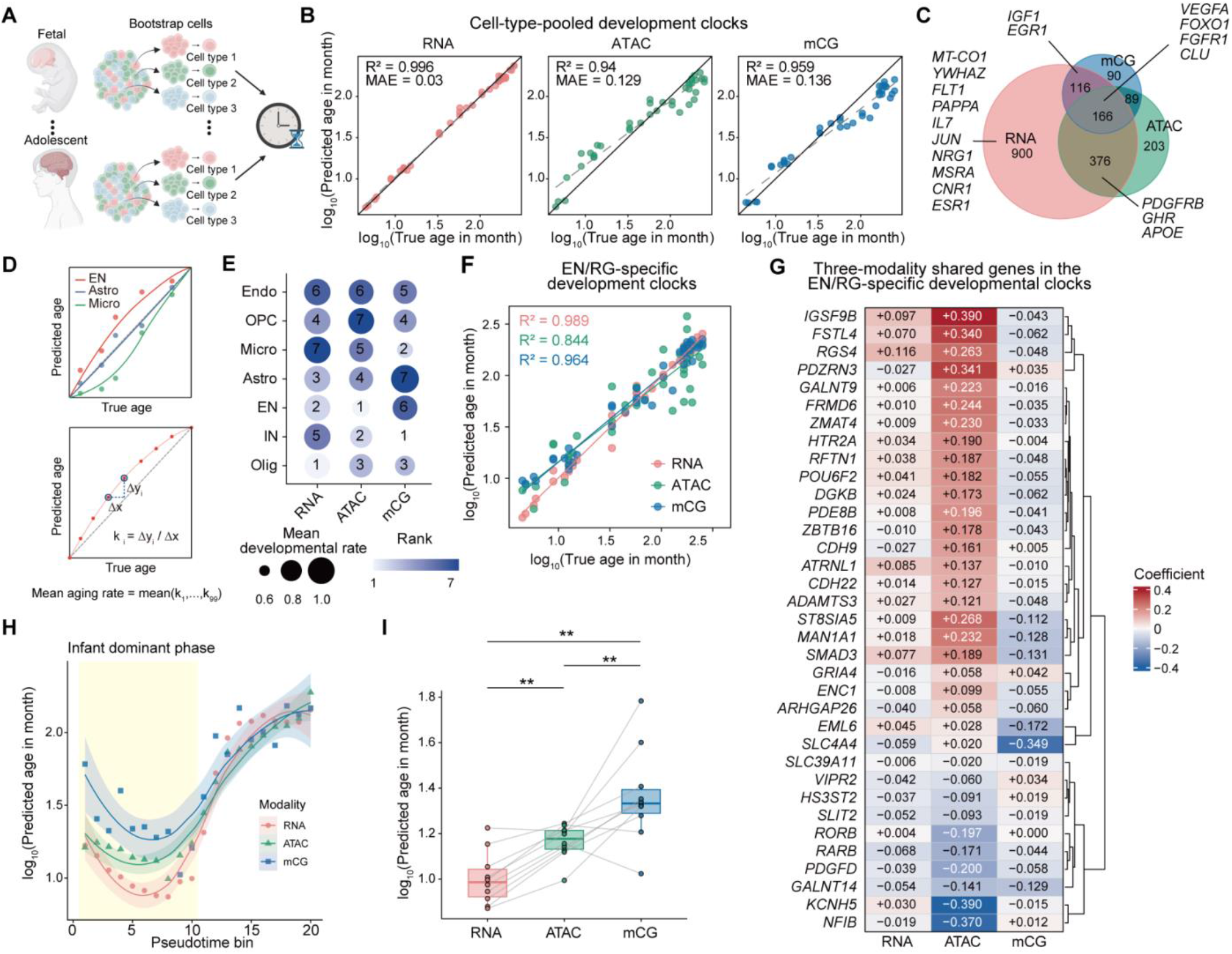
Modality-specific developmental clocks reveal distinct temporal representations of brain maturation. **(A)** Schematic of the developmental clock framework. Single cells were aggregated into bootstrap pseudocells by donor and cell type. **(B)** Cell-type-pooled developmental clock performance in RNA, ATAC, and mCG. Each point represents one donor, with predicted log-age summarized as the median across pseudocells. Dashed line: linear fit; solid line: identity (y = x). **(C)** Venn diagram of model-contributing genes across modalities. For ATAC and mCG models, tile-level features were mapped to their linked genes. Selected neurodevelopmental or aging-related genes are labeled. **(D)** Schematic of developmental rate calculation. For each modality and cell type, smoothing splines were fitted to median predicted versus true log-age. Developmental rate was defined as the mean slope across 99 intervals between 100 equally spaced fitted points. **(E)** Cell-type-specific developmental rate rankings across modalities. Dot size indicates mean developmental rate. Dot color indicates modality-specific rank. Cell types are ordered by summed ranks. **(F)** EN/RG-specific developmental clock performance for the three modalities. Solid line: linear fit. **(G)** Heatmap of LASSO coefficients for genes shared across all three EN/RG-specific developmental clocks. **(H)** Modality-specific predicted developmental ages along the EN L2/3 IT pseudotime trajectory. Lines show LOESS-smoothed trends with 95% CI. Yellow shading highlights early pseudotime bins (1–10). **(I)** Paired comparison of predicted log-ages across modalities for early pseudotime bins (bins 1–10). Lines connect matched bins. Paired Wilcoxon signed-rank tests with Benjamini–Hochberg correction. **adjusted *p* < 0.01.

We next examined how developmental progression varied across cell types. Modality-specific clocks revealed substantial variations in relative developmental rate (**Fig. 5D, Fig. S7E, F**), with endothelial/vascular cells, OPCs, and microglia consistently ranking among the fastest-progressing populations (**Fig. 5E**). Notably, ENs ranked among the slowest populations in accessibility and gene expression but among the fastest in mCG. Cell-type-specific models further identified predominantly lineage-restricted age-associated features linked to neuronal development, myelination, and immune regulation (**Fig. S7E–H**). These observations suggest that different regulatory layers capture distinct aspects of cell-type-specific maturation.

To directly compare the developmental timing within the same lineage, we next constructed EN-specific clocks, which accurately predicted ages across all three modalities (RNA R^2^ = 0.989, ATAC R^2^ = 0.844, mCG R^2^ = 0.964) (**Fig. 5F, G, Fig. S7I, J)**. Although shared features generally reflected coordinated transcriptional activation, chromatin opening, and DNA demethylation, the clocks assigned systematically different ages along the same L2/3 IT trajectory (**Fig. 5H**). During the early postnatal maturation (which we referred to as the infant-dominated phase), mCG-based predictions were significantly more advanced than both ATAC- and RNA-based predictions, with ATAC also preceding RNA (**Fig. 5I**, paired Wilcoxon signed-rank tests, BH-adjusted *p* < 0.01). The mCG-associated temporal offset remained significant across the full trajectory (**Fig. S7K, L**), suggesting that DNA methylation captures developmental states not yet fully reflected in transcriptional or accessibility profiles.

Together, these findings reveal that developmental age is represented differently across molecular modalities and cellular lineages. In conjunction with prolonged methylation remodeling, the advanced mCG-derived developmental states suggest that DNA methylation encodes a distinct temporal dimension of human brain maturation, consistent with epigenetic priming in which methylation remodeling marks regulatory states before corresponding accessibility and transcriptional programs are fully established.

### Complementary epigenomic networks converge on shared regulators governing excitatory neuron maturation

Having established a temporal hierarchy of epigenomic remodeling, we next asked whether these two epigenomic layers converge on common transcriptional regulators during excitatory neuron maturation. We applied the FigR framework ^29^ independently to ATAC–RNA and mCG–RNA profiles along the EN L2/3 IT developmental trajectory, integrating transcription factor (TF) motif enrichment with TF expression and associated DORC activity to infer modality-specific gene regulatory networks (GRNs, **Fig. S8**, **Table S15, S16**).

Despite substantial differences in their inferred regulatory interactions, the two regulatory networks converged on core developmental TFs, particularly EGR1 and EGR3 (**Fig. 6A, B**). These activity-dependent regulators ^30,31^ showed opposite regulation-score directions across modalities, reflecting the expected relationship between transcription activation, chromatin opening, and demethylation at their associated regulatory regions. Beyond this shared core, the ATAC-based network highlighted maturation-associated TFs such as NR4A1 and PPARG ^32,33^, together with progenitor-associated TFs, including LHX2, EBF4, TCF12, and PAX6 ^34–36^, whereas the mCG-based network identified a distinct set of TFs including BACH1, BACH2, FOXG1, KLF9, and NR3C2. Integration of both networks revealed a broader regulatory architecture connecting shared and modality-specific TFs to developmental DORC genes (**Fig. 6C**). The developmental patterns of selected core TFs further showed coordinated changes in RNA expression, accessibility-associated motif activity, and methylation-associated motif score (**Fig. 6D**), suggesting that common transcriptional programs are represented through distinct epigenomic regulatory relationships.

**Figure 6.**
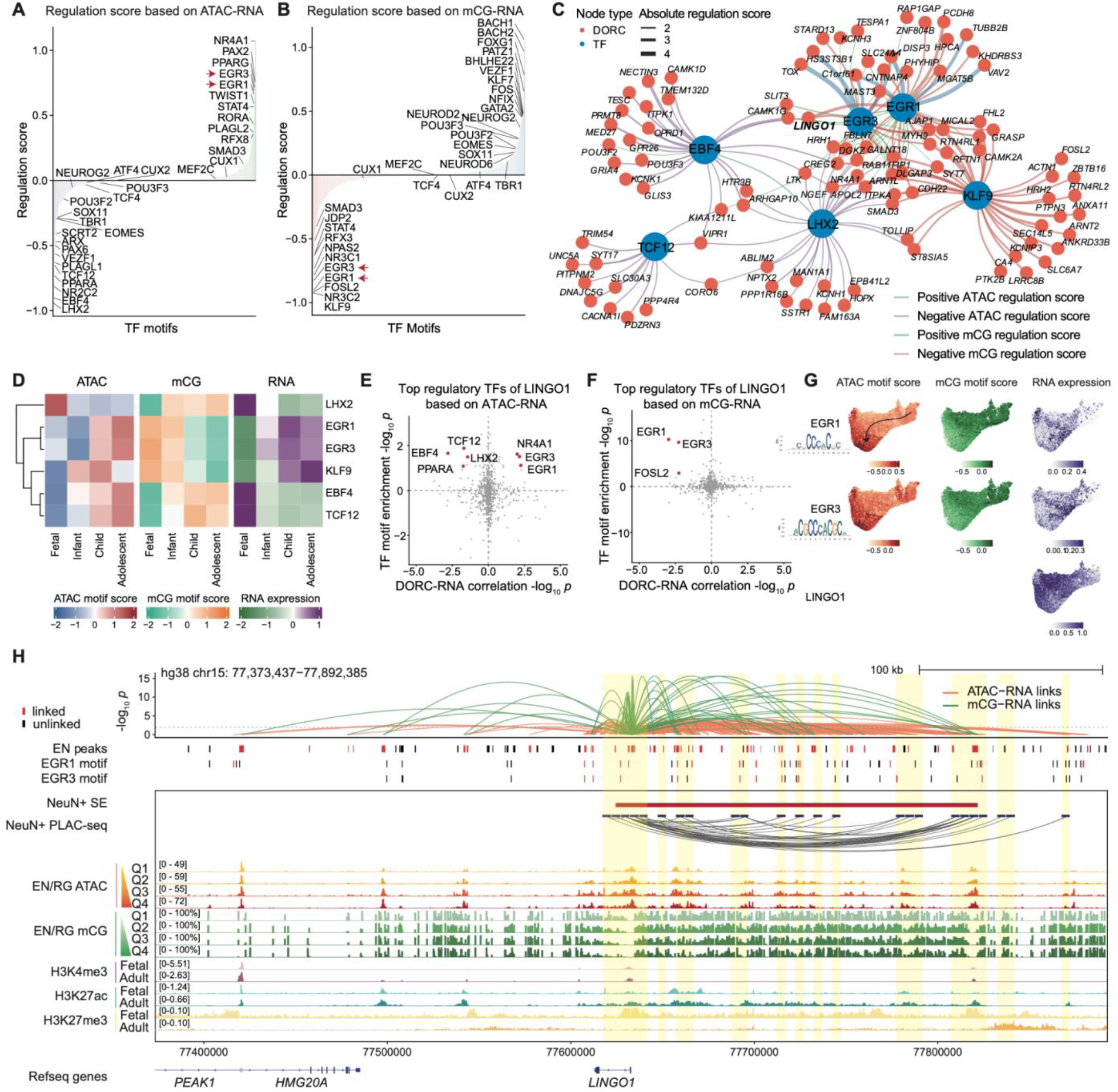
Multimodal gene regulatory networks converge on shared core TFs governing excitatory neuron maturation. **(A)** Ranked mean TF regulation scores based on ATAC–RNA FigR analysis along the EN L2/3 IT pseudotime trajectory. Each bar represents one TF. Positive and negative scores indicate corresponding correlations between TF RNA expression and target DORC score. Top-ranked positive and negative TFs and selected cortical developmental TFs are labeled. **(B)** As in (A), for mCG–RNA regulatory analysis. Positive and negative scores indicate corresponding correlations between TF RNA expression and target mDORC score. **(C)** Merged TF–DORC regulatory network integrating ATAC-based and mCG-based results. TF nodes (blue) are sized by the number of regulated DORCs. DORC nodes (coral) represent target genes. Edge width is proportional to the absolute regulation score. **(D)** Developmental stage-resolved heatmap of ATAC motif scores, mCG motif scores, and RNA expression for six core TFs from the merged network. Values are *z*-score standardized across stages within each TF. **(E)** Top TFs regulating *LINGO1* identified by ATAC–RNA regulatory analysis. Each point represents a TF–*LINGO1* pair. Top TFs are labeled. **(F)** As in (E), for mCG–RNA regulatory analysis. **(G)** UMAP of ATAC motif scores, mCG motif scores, and RNA expression for EGR1, EGR3, and LINGO1 along the EN L2/3 IT pseudotime trajectory. Arrow: early-to-late trajectory. **(H)** Genome tracks around the *LINGO1* locus. Tracks from top to bottom: ATAC–RNA (orange) and mCG–RNA (green) tile–gene link significance; EN ATAC peaks, EGR1 and EGR3 motif positions colored red if overlapping a linked tile and black otherwise; NeuN+ SE annotations; NeuN+ PLAC-seq chromatin loops ^15^ highlighted by yellow areas; ATAC signal and mCG fraction across four pseudotime quartile bins (Q1–Q4) of the EN L2/3 IT lineage; histone modification signals in fetal and adult brain.

This pattern is particularly notable given the established role of EGR1 in recruiting the DNA demethylase TET1 to shape the brain methylome during postnatal development ^30^, and the recent finding that EGR1 binding at enhancers is enriched in excitatory neurons in a subtype-specific manner ^37^. In contrast, early developmental regulators TCF12 and LHX2 showed the opposite trajectory, consistent with their roles in progenitor maintenance and early cortical development ^35,38^. These findings indicate that common developmental TF programs can be represented through distinct accessibility- and methylation-defined regulatory architectures.

To examine this convergence at individual loci, we focused on *LINGO1*, a neuronal regulator associated with neurite outgrowth and maturation ^39^. *LINGO1* was identified as a prominent DORC in both modalities (**Fig. S2H, I**) and linked to multiple TFs, including EGR1 and EGR3 (**Fig. 6E, F**). Their motif activities varied with *LINGO1* RNA expression along the excitatory neuron trajectory (**Fig. 6G**), consistent with coordinated regulation of this locus during neuronal maturation. At the genomic level, ATAC–RNA and mCG–RNA regulatory links overlapped predicted EGR-binding motifs within an extended upstream regulatory domain that also intersected neuronal super-enhancers and PLAC-seq chromatin interactions (**Fig. 6H**) ^15^. This locus also showed a developmental transition from fetal H3K27me3 enrichment towards adult H3K27ac enrichment, consistent with progressive activation ^40^. Genome-wide, beyond the *LINGO1* locus, linked elements from both modalities were significantly enriched within PLAC-seq interaction domains of NeuN-positive super-enhancers (SEs; 1.28-fold for mCG, *p* = 2.7e-14; 1.23-fold for ATAC, *p* = 1.4e-32), further supporting that both ATAC- and methylation-based *cis*-regulatory mapping may precisely delineate physically interacting regulatory elements.

Together, these analyses reveal that chromatin accessibility and DNA methylation capture complementary TF–target architectures while converging on shared developmental regulators, providing a more integrated view of the regulatory programs underlying neuron maturation.

### Developmental epigenomic programs capture complementary neuropsychiatric genetic risk

Having characterized the distinct regulatory architectures and developmental dynamics of chromatin accessibility and DNA methylation, we next investigated how these complementary epigenomic landscapes inform the genetic basis of neuropsychiatric disorders. We leveraged cell type-resolved putative regulatory links and GRNs to place non-coding disease variants within regulatory circuits connecting *cis*-regulatory elements, candidate target genes, and upstream transcription factors.

Stratified LD score regression (S-LDSC) revealed significant enrichment of schizophrenia (SCZ), bipolar disorder (BD), and major depressive disorder (MDD) heritability in EN-linked regulatory elements defined by both accessibility and mCG (**Fig. S9A**). When partitioning these elements into ATAC-specific, mCG-specific, and shared classes, all three classes showed significant heritability enrichment (**Fig. 7A**), highlighting complementary disease-relevant regulatory information captured through chromatin accessibility and DNA methylation.

**Figure 7.**
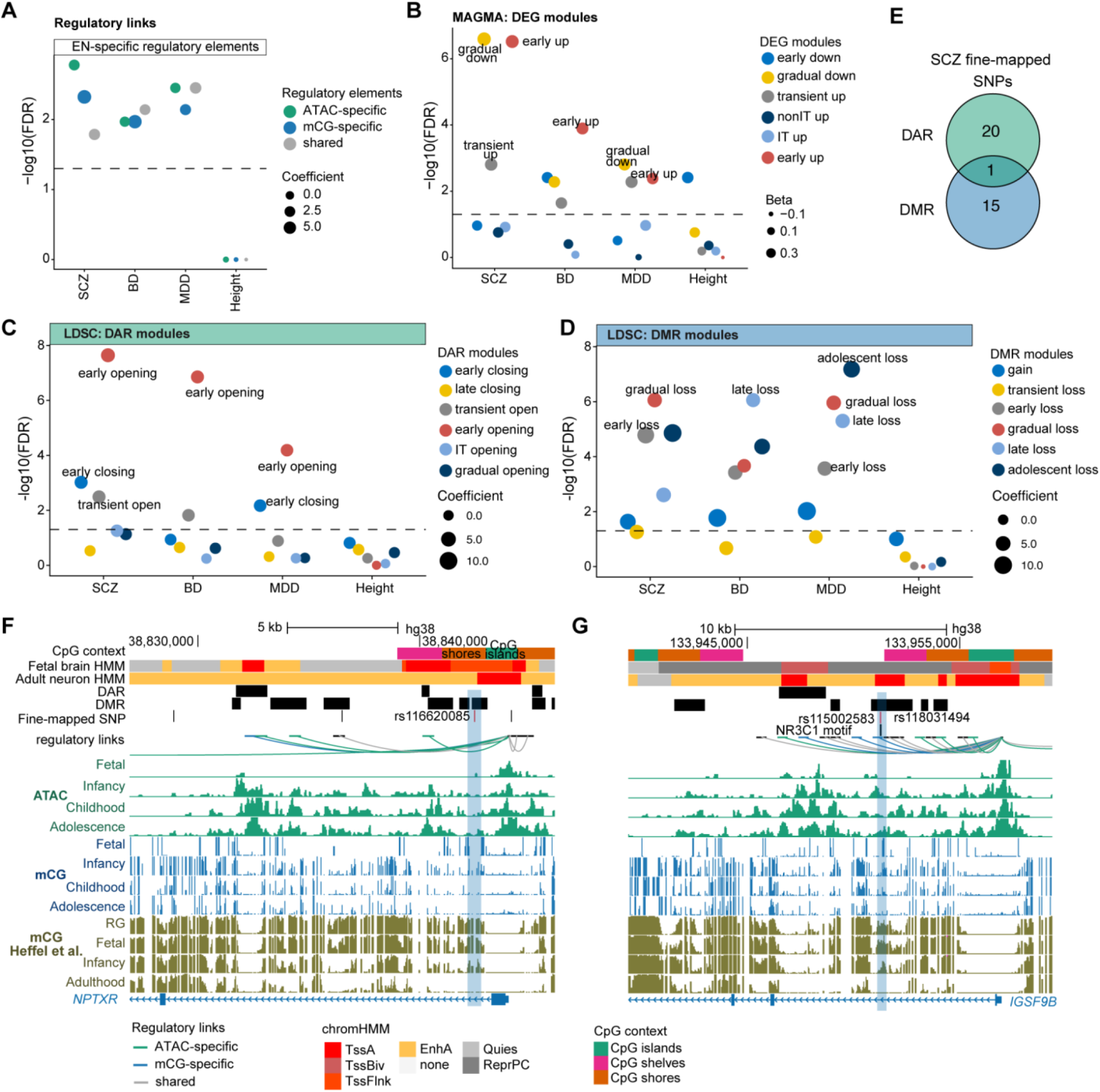
Developmental epigenomic programs capture complementary neuropsychiatric genetic risk. **(A)** Stratified linkage disequilibrium score regression (S-LDSC) analysis of heritability enrichment for schizophrenia (SCZ), bipolar disorder (BD), and major depressive disorder (MDD) in EN-specific regulatory elements classified as ATAC-specific, mCG-specific, or shared between modalities, with height included as a control trait. **(B)** MAGMA gene-set associations of SCZ, BD, MDD, and height with DEG modules across EN development. **(C)** S-LDSC analysis of heritability enrichment of neuropsychiatric disorders across developmental DAR modules. **(D)** S-LDSC analysis of heritability enrichment of neuropsychiatric disorders across developmental DMR modules. In (A–D), dashed horizontal lines indicate the significance threshold (*FDR* = 0.05). Point sizes represent normalized S-LDSC coefficients in (A, C, D) and MAGMA beta coefficients in (B). **(E)** Venn diagram showing the number of SCZ fine-mapped SNPs (PIP > 0.5) overlapping developmental DMRs and DARs during EN maturation. **(F)** Genomic tracks of the *NPTXR* locus highlighting the fine-mapped SCZ variant rs116620085 (PIP = 0.9993) overlapping a developmental DMR located within a CpG shore. Tracks show CpG context, fetal and adult neuronal chromatin states, DARs, DMRs, fine-mapped SCZ variants, tile–gene links, and developmental ATAC and mCG profiles. Independent mCG profiles from Heffel et al. are shown for validation ^8^. **(G)** Genomic tracks of the *IGSF9B* locus highlighting fine-mapped SCZ variants (rs115002583 and rs118031494; PIP = 1) intersecting an mCG-specific tile–gene element linked to the DORC gene *IGSF9B*.

We next asked whether neuropsychiatric genetic risk was preferentially localized to developmental programs remodeled during EN maturation. Developmental DARs and DMRs, but not the gene bodies of DEGs, showed significant heritability enrichment (**Fig. S9B**), implicating epigenomically remodeled regions in neuropsychiatric genetic risk. Genetic risk for SCZ, BD, and MDD was significantly associated with multiple developmental transcriptional programs and enriched in epigenomically remodeled regions, particularly transcriptionally upregulated genes, chromatin-opening regions, and methylation-loss regions (**Fig. 7B–D, Fig. S9B**). Despite this shared association with activation-related programs, their temporal patterns differed across modalities: transcriptional and chromatin accessibility associations were most pronounced in early upregulation and early chromatin opening, respectively, whereas DMRs showed substantial enrichment across early, gradual, late, and adolescent methylation-loss programs (**Fig. 7B–D**), highlighting the prolonged developmental involvement of DNA methylation remodeling in neuropsychiatric genetic risk. Enrichment in DMRs also extended to CpG shores and shelves that are less prominently represented by accessibility changes (**Fig. S9C**). Conditional S-LDSC analyses further revealed significant enrichment for both DARs and DMRs when jointly modeled (**Fig. S9D**), supporting complementary disease-relevant regulatory information captured by chromatin accessibility and DNA methylation.

We next integrated SCZ fine-mapping ^41^ with developmental regulatory annotations to resolve these associations at individual loci. Among fine-mapped SCZ variants with posterior inclusion probability (PIP) > 0.5, 15 specifically overlapped DMRs and 20 specifically overlapped DARs, with only one overlapping both (**Fig. 7E**, **Table S17**), further supporting the complementary regulatory information provided by the two epigenomic modalities. For example, at the *NPTXR* locus, rs116620085 (PIP = 0.9993) fell within a DMR-specific CpG-shore region (**Fig. 7F**), consistent with the enrichment of disease-associated DMRs in CpG-flanking regions. At the *IGSF9B* locus, fine-mapped variants with PIP = 1 (rs115002583 and rs118031494) intersected an mCG-specific regulatory element linked to *IGSF9B*, a DORC gene with regulatory associations enriched for NR3C1 (**Fig. 7G**). Variants fell within a predicted NR3C1 binding motif, concordant with the GRN-inferred regulatory context at this locus.

Together, these analyses show that chromatin accessibility and DNA methylation capture complementary components of neuropsychiatric genetic risk, connecting disease-associated non-coding variants to candidate regulatory elements, developmental genes, and upstream transcription factors.

## Discussion

Human brain maturation involves profound changes in gene expression and epigenetic regulation, yet the extent to which these molecular layers encode shared regulatory information and follow coordinated developmental trajectories has remained unclear. By jointly profiling transcription, chromatin accessibility, and DNA methylation across human brain development, we reveal two important principles: regulatory information is distributed across partially distinct epigenomic architectures, and these architectures remodel over different developmental timescales. These findings extend prior observations of temporally distinct epigenomic remodeling ^8^ and reveal complementary dimensions of cellular development.

An important implication of our findings is that chromatin accessibility alone does not capture all of the *cis*-regulatory associations detected by the combined analysis. Although accessibility and DNA methylation converge on many canonical developmental regulators, they identify substantially different *cis*-regulatory associations and TF–target relationships. This distinction suggests that the two modalities capture different aspects of regulatory organization rather than merely providing alternative measurements of the same regulatory elements. Importantly, methylation-associated regulatory information is strongly influenced by CpG context ^42–44^, highlighting genomic features that may not be adequately represented by conventional accessibility-based annotations. Together, these observations argue for a more integrated view of *cis*-regulatory architecture in which complementary epigenetic measurements provide a more complete view of developmental gene regulation.

The prolonged remodeling of DNA methylation raises a broader question about the relationship between cellular identity and epigenetic maturation. Although transcriptional programs undergo major changes during neuronal differentiation, the associated methylation landscape continues to evolve long after these transitions ^45,46^. This temporal separation suggests that the acquisition of transcriptional identity does not necessarily mark the completion of molecular maturation. Instead, DNA methylation may capture a slower dimension of development, progressively reflecting regulatory refinement or the accumulated history of cellular differentiation ^9,47^. The distinct developmental states inferred from modality-specific clocks further suggest that molecular maturation cannot be represented by a single, unified trajectory. Together, these observations point toward a multilayered model of brain development in which transcriptional identity and epigenetic maturation proceed on partially independent timescales, potentially allowing cells with similar transcriptional states to retain distinct developmental epigenetic signatures. Thus, DNA methylation provides not only complementary regulatory information but also a distinct temporal perspective on molecular maturation.

These findings also suggest a broader framework for interpreting the genetic basis of neuropsychiatric disorders. If regulatory information is distributed across distinct epigenetic layers that remodel on different timescales, disease-associated variation may influence developmental programs not fully captured by transcriptional or accessibility-based annotations ^48^. The enrichment of neuropsychiatric genetic risk across both early and prolonged methylation-loss programs underscores the importance of considering the developmental timing of disease-associated regulatory remodeling. Rather than treating the epigenome as a static annotation of genetic variation, our findings support a developmental perspective in which regulatory landscapes continuously evolve as the brain matures. The prolonged remodeling of neuronal DNA methylation raises the possibility that susceptibility to genetic and environmental perturbations may extend across a broader developmental period than previously appreciated ^8,49,50^. Integrating complementary regulatory modalities across developmental time may therefore provide a more complete foundation for understanding how non-coding genetic variation intersects with human brain development and neuropsychiatric disease susceptibility.

### Limitations of the study

Several limitations should be considered. First, the sparsity of single-cell epigenomic measurements necessitated aggregation for certain analyses. Moreover, ME-seq preferentially captures DNA methylation at accessible regulatory regions, limiting comprehensive assessment of genome-wide methylation dynamics, including methylation within gene bodies and non-CG methylation (mCH). Second, uneven developmental sampling, particularly the limited representation of progenitors and immature neurons, may constrain the resolution of early developmental transitions. Third, the cross-sectional nature of our dataset precludes direct inference of causal or temporal relationships among regulatory layers. Although modality-specific developmental clocks reveal systematic differences in predicted developmental age, these offsets do not establish the molecular ordering of regulatory events. Furthermore, ME-seq does not distinguish 5mC from 5hmC, which may contribute to the observed methylation dynamics. Future studies incorporating longitudinal or experimentally tractable developmental models and more comprehensive methylation profiling will be important for resolving these questions.

## Resource Availability

## Lead contact

Sai Ma

## Materials availability

All unique/stable reagents generated in this study are available from the Lead Contact with a completed Materials Transfer Agreement.

## Acknowledgments

We thank all the members of the Ma and Roussos labs for critical reading of the manuscript and helpful discussions. This work was supported by the National Institutes of Health (NIH), including the National Institute of Mental Health (NIMH) under awards R01-MH110921 (to P.R.), U01-MH116442 (to P.R.), and R01-MH125246 (to P.R.); the National Institute on Aging (NIA) under awards R21-AG092994 (to S.M.), R01-AG050986 (to P.R.), R01-AG065582 (to P.R.), R01-AG067025 (to P.R.), R01-AG082185 (to P.R. and D.L.), U24-AG087563 (to P.R.), and R01-AG095776 (to P.R.); and the National Cancer Institute (NCI) under awards R61-CA297881 (to S.M.) and U01-CA290442 (to S.M.). All research involving human fetal tissue from elective abortions was completed before 22 January 2026, the effective date of NIH Guide Notice NOT-OD-26-028. No NIH funds were used for research involving such tissue after that date, and subsequent work was limited to analysis of existing data and preparation of this manuscript.

## Author contributions

S.M. and P.R. conceived and supervised the research. B.Z. and Z.L. generated the single-cell sequencing library. N.T. provided human brain samples. F.A.D. and D.A.K. provided organoid samples. L.S., Z.W., S.M., and C.W. conducted the data analysis. S.M., P.D., P.R., J.F.F., D.L., and K.G. supervised data analyses. S.M., L.S., and Z.W. wrote the manuscript, and P.D. and P.R. revised the manuscript. All authors approved the manuscript.

## Declaration of generative AI and AI-assisted technologies in the manuscript preparation process

During the preparation of this work, the authors used ChatGPT to assist with language editing and improving the clarity of the manuscript. The authors reviewed and edited the content as needed and take full responsibility for the content of the published article.

## Supplemental information

Supplementary figures. Figures S1–S9

Supplementary tables. Tables S1–S17

## Supplementary Tables

**Table S1.** Summary of human brain samples

**Table S2.** Cell-type-specific differentially methylated regions

**Table S3.** Significant ATAC-RNA tile-gene links

**Table S4.** Significant mCG-RNA tile-gene links

**Table S5.** Functional enrichment of ATAC-specific DORC genes

**Table S6.** Functional enrichment of mCG-specific DORC genes

**Table S7.** Pseudotime along EN development across three molecular modalities

**Table S8.** Differentially expressed genes across EN maturation

**Table S9.** Differentially accessible regions across EN maturation

**Table S10.** Differentially methylated regions across EN maturation

**Table S11.** Pseudotime along oligodendroglial development across three modalities

**Table S12.** Differentially expressed genes across oligodendroglial maturation

**Table S13.** Differentially accessible regions across oligodendroglial maturation

**Table S14.** Differentially methylated regions across oligodendroglial maturation

**Table S15.** Global FigR regulatory profile based on ATAC–RNA within the EN L2/3 IT lineage

**Table S16.** Global FigR regulatory profile based on mCG–RNA within the EN L2/3 IT lineage

**Table S17.** Fine-mapped SNPs overlapping with DMRs or DARs during EN maturation

## STAR Methods

### Key Resources Table

| REAGENT or RESOURCE | SOURCE | IDENTIFIER |
|---|---|---|
| **Bacterial and Virus Strains** |  |  |
| pcDNA3-Flag-Tet1 CD | Addgene | 70129 |
| **Chemicals, Peptides, and Recombinant Proteins** |  |  |
| RNase Inhibitor | QIAGEN | Y9240L |
| SUPERase·In RNase Inhibitor | Thermo Fisher Scientific | AM2696 |
| 60% Iodixanol | Sigma-Aldrich | D1556-250ML |
| 1M Tris HCl pH 7.5 | Thermo Fisher Scientific | 15567027 |
| 1M Tris HCl pH 8.0 | Thermo Fisher Scientific | 15568025 |
| Igepal 630 | Sigma-Aldrich | I8896-50ML |
| cOmplete Protease Inhibitor | Sigma-Aldrich | 11697498001 |
| 16% Formaldehyde (w/v) | Thermo Fisher Scientific | 28906 |
| Tagmentase | Diagenode | C01070010-100 |
| Tris-acetate buffer | Bioworld | 40120265-2 |
| Dimethylformamide | Thermo Fisher Scientific | 20673 |
| Deoxynucleotide Solution Mix | NEB | N0447L |
| 100 mM GTP | Thermo Fisher Scientific | R1461 |
| Dithiothreitol | Thermo Fisher Scientific | 70726-5ML |
| PEG 8000 | Thermo Fisher Scientific | BP233-100 |
| Maxima H Minus Reverse Transcriptase | Thermo Fisher Scientific | EP0753 |
| T4 DNA ligase | NEB | M0202L |
| Proteinase K | Sigma-Aldrich | P2308 |
| Dynabeads MyOne Streptavidin C1 | Thermo Fisher Scientific | 65001 |
| Ficoll solution | Sigma-Aldrich | F5415-25ML |
| NxGen RNase Inhibitor | Lucigen | 30281-2 |
| KAPA HiFi HotStart ReadyMix | Fisher Scientific | NC0295239 |
| Biotium EvaGreen 20X | Fisher Scientific | NC0521178 |
| AMPure XP | Beckman Coulter | A63880 |
| NEBNext High-Fidelity 2X PCR Master Mix | NEB | M0541L |
| T4 DNA polymerase | Thermo Fisher Scientific | EP0062 |
| Ampligase Thermostable DNA Ligase | LGC Genomics | A0110K |
| Adenosine 5'-Triphosphate | NEB | P0756S |
| NEBNext Enzymatic Methyl-seq Conversion Module | NEB | E7125S |
| Pyridine borane | Fisher Scientific | AAL1317809 |
| DNA Clean & Concentrator-5 | Zymo | D4014 |
| KAPA HiFi HotStart Uracil+ ReadyMix Kit | Fisher Scientific | 50-196-5287 |
| SYBR Green I Nucleic Acid Gel Stain | Thermo Fisher Scientific | S7563 |
| **Deposited Data** |  |  |
| Human brain ME-seq data | This manuscript | In progress |
| Organoid data | This manuscript | In progress |
| snm3C-seq3 data | Heffel et al. <sup>8</sup> | GSE213950 |
| PLAC-seq data | Nott et al. <sup>15</sup> | phs001373.v2.p2 |
| Super-enhancer data | Hnisz et al. <sup>22</sup> | GSE51522 |
| ChromHMM data | Rahman et al. <sup>23</sup> | syn26148119 |
| External validation dataset 1 | Zhu et al. <sup>27</sup> | GSE204684 |
| External validation dataset 2 | Fullard et al. <sup>28</sup> | RRID:SCR_021059 |
| External validation dataset 3 | Wang et al. <sup>13</sup> | nemo:dat-oiif74w |
| Histone modification data | Dong et al. <sup>40</sup> | syn25672226 |
| **Software and Algorithms** |  |  |
| R (v4.3.2) |  | https://www.R-project.org |
| STAR (v2.7.5) |  | https://github.com/alexdobin/STAR |
| Bismark |  | https://github.com/FelixKrueger/Bismark |
| MACS2 |  | https://github.com/macs3-project/MACS |
| featurecounts |  | http://subread.sourceforge.net |
| Seurat (v5.0.0)<sup>51</sup> |  | https://github.com/satijalab/seurat/ |
| SoupX(v1.6.2)<sup>52</sup> |  | https://github.com/constantAmateur/SoupX |
| Scanpy (v1.10.1)<sup>53</sup> |  | https://github.com/scverse/scanpy |
| SnapATAC2 (v2.8.0)<sup>54</sup> |  | https://github.com/scverse/SnapATAC2 |
| BBKNN (v1.6.0)<sup>55</sup> |  | https://github.com/teichlab/bbknn |
| Signac (v1.10.1)<sup>56,57</sup> |  | https://stuartlab.org/signac/ |
| chromVAR (v1.26.0)<sup>58</sup> |  | https://github.com/GreenleafLab/chromVAR |
| Slingshot (v2.10.0)<sup>59</sup> |  | https://github.com/kstreet13/slingshot |
| tradeSeq (v1.12.0)<sup>60</sup> |  | https://github.com/statomics/tradeSeq |
| scVI (v1.3.0)<sup>61</sup> |  | https://github.com/scverse/scvi-tools |
| LDSC (v1.0.0)<sup>62</sup> |  | https://github.com/bulik/LDSC |
| MAGMA (v.1.07b)<sup>63</sup> |  | https://ctg.cncr.nl/software/magma |
| FigR (v0.1.0)<sup>29</sup> |  | https://buenrostrolab.github.io/FigR/ |
| EnrichR (v3.4)<sup>64</sup> |  | https://github.com/guokai8/Enrichr/ |
| rGREAT (v2.9.2)<sup>65</sup> |  | https://github.com/jokergoo/rGREAT |
| edgeR (v4.0.16)<sup>66</sup> |  | https://bioconductor.org/packages/release/bioc/html/edgeR.html |
| gprofiler2 (v0.2.3)<sup>67</sup> |  | https://github.com/egonw/r-gprofiler2 |
| ChIPseeker (v1.38.0)<sup>68</sup> |  | https://github.com/YuLab-SMU/ChIPseeker |
| GenomicScores (v2.25.1)<sup>69</sup> |  | https://github.com/rcastelo/GenomicScores |
| topGO (v2.65.0)<sup>70</sup> |  | https://github.com/federicomarini/topgo |
| glmnet (v5.0)<sup>71</sup> |  | https://github.com/cran/glmnet |
| motifmatchr (1.34.0) |  | https://github.com/GreenleafLab/motifmatchr/ |
| BPCells (v0.3.1) |  | https://github.com/bnprks/BPCells |

### Sample collection

Postmortem human brain specimens were obtained from 40 donors spanning fetal development through adolescence, ranging from 18 gestational weeks (gw) to 20 years of age. Fetal specimens were collected from de-identified prenatal autopsies without neuropathological abnormalities at the Icahn School of Medicine at Mount Sinai, where the cortical plate was freshly dissected from the anterior frontal lobe. Postnatal samples (birth to 20 years) of the dorsolateral prefrontal cortex (DLPFC) were accessed from the Human Brain Collection Core (HBCC), the University of Miami Brain Endowment Bank, and the Mount Sinai Brain Bank.

Donors were grouped into four developmental stages, including fetal (18–24 gw; n = 4), infancy (0–1 year; n = 8), childhood (2–10 years; n = 12), and adolescence (11–20 years; n = 16). Specimens from male and female donors were included across each developmental stage. All tissue collection and research procedures were approved by the respective Institutional Review Boards. Detailed donor characteristics, including age, sex, brain region, and tissue source, are provided in **Table S1**.

### Cortical organoid differentiation

Human pluripotent stem cells (hPSCs), specifically the H9 embryonic stem cell line and MSN25 induced pluripotent stem cell line, were dissociated into cell aggregates using phosphate-buffered saline (PBS) containing EDTA and transferred to ultra-low-attachment six-well plates containing neural induction medium (NIM) supplemented with the CEPT cytoprotective cocktail. The day of aggregation was designated day 0. NIM consisted of Glasgow’s modified Eagle’s medium (GMEM; 384 mL), knockout serum replacement (100 mL), antibiotic–antimycotic (5 mL), non-essential amino acids (5 mL), and sodium pyruvate (5 mL). Freshly prepared NIM was supplemented with 0.1 mM β-mercaptoethanol, 10 μM SB431542, 250 nM LDN-193189 (MedChemExpress), 3 μM IWR1-endo, 1× CEPT cytoprotective cocktail and 1× polyamine supplement. The CEPT cocktail consisted of 50 nM chroman 1, 5 μM emricasan, 0.7 μM trans-integrated stress response inhibitor (trans-ISRIB) and polyamine supplement, prepared at the concentrations specified above. NIM was completely replaced on day 3 with fresh NIM containing CEPT and on day 6 with NIM without CEPT.

On day 9, organoids were transferred to an orbital shaker for suspension culture. From days 9–35, organoids were maintained in Maintenance Medium 1, consisting of a 1:1 mixture of DMEM/F12 and Neurobasal medium, supplemented with B27 without vitamin A (2%), N2 (1%), non-essential amino acids (1%), GlutaMAX (1%) and antibiotic–antimycotic (1%), supplemented with 50 μM β-mercaptoethanol. FGF2 and EGF were added at 10 ng/mL each from days 9–25. Chemically defined lipids were added at 1× concentration beginning on day 18. From day 25 onward, FGF2 and EGF were omitted, and BDNF and NT-3 were added at 10 ng/mL each.

From day 36, organoids were maintained in Maintenance Medium 2, consisting of a 1:1 mixture of DMEM/F12 and Neurobasal medium, supplemented with B27 with vitamin A (2%), N2 (1%), GlutaMAX (1%), chemically defined lipids (1%) and antibiotic–antimycotic (1%). The medium was supplemented with 50 μM β-mercaptoethanol, 2 nM ascorbic acid, 1% Matrigel, 5 μg/mL heparin, 10 ng/mL BDNF and 10 ng/mL NT-3. From days 56–62, leukemia inhibitory factor (LIF) was additionally included at 10 ng/mL. LIF supplementation was discontinued after day 62, and organoids were maintained in Maintenance Medium 2 through day 75. Complete medium exchanges were performed every 2–3 days throughout the culture period, with additional changes performed as needed.

### Nuclei isolation from prefrontal cortex

Human brain single-nucleus suspensions were prepared following the OMNI-ATAC ^72^ protocol with minor modifications. Frozen tissue was placed into a pre-chilled 2 mL Dounce homogenizer containing 450 μL of 1× homogenization buffer (320 mM sucrose, 0.1 mM EDTA, 0.1% NP-40, 5 mM CaCl_2_, 3 mM Mg(Ac)_2_, 10 mM Tris-HCl, pH 8.0, 16 μM PMSF, 0.15 mM DTT, and 0.26% Enzymatics RNase Inhibitor, Y9240L, Qiagen). The tissue was homogenized using 5 strokes with pestle A, followed by 15 strokes with pestle B. Next, 400 μL of the homogenate was transferred to a pre-chilled 2 mL graduated microcentrifuge tube. An equal volume (400 μL) of 50% iodixanol solution (prepared in 1× homogenization buffer) was added and mixed to create a 25% iodixanol layer. Subsequently, 600 μL of 29% iodixanol solution (160 mM sucrose in 1× homogenization buffer) was carefully layered beneath the 25% iodixanol layer, followed by 600 μL of 35% iodixanol solution (160 mM sucrose in 1× homogenization buffer) layered at the bottom. Nuclei were isolated by centrifugation at 3,000 g for 20 minutes at 4°C using a swinging-bucket rotor. The resulting nuclear pellet was resuspended in HCBI buffer (0.26% Enzymatics RNase Inhibitor, 0.04% BSA, 10 mM NaCl, 3 mM MgCl_2_, and 10 mM HEPES, pH 7.2) and immediately processed for fixation.

### Library construction and ME-seq

ME-seq libraries were prepared as described in an online protocol ^73^. Briefly, nuclei were fixed with 0.2% formaldehyde and transposed with Tn5 transposase to mark regions of open chromatin. The mRNA was reverse transcribed using a poly(T) primer containing a unique molecular identifier (UMI) and a random primer. Both primers contained a biotin tag. Permeabilized cells were distributed in a 96-well plate to hybridize well-specific barcoded oligonucleotides to transposed chromatin fragments and poly(T) cDNA. Hybridization was repeated three times to expand the barcoding space and ligate cell barcodes to cDNA and chromatin fragments. Reverse crosslinking was performed to release barcoded molecules. cDNA was separated from chromatin using streptavidin beads, and each library was prepared separately for sequencing. Chromatin DNA was purified, gap-filled and oxidized using mTET1, and deaminated with 1 M pyridine borane before amplification. Both DNA and RNA libraries were quantified with KAPA Library Quantification Kit and pooled to 4 nM for sequencing. Libraries were sequenced on the Illumina NovaSeq platform with 150-bp paired-end reads (Read 1: 150 cycles, Index 1: 8 cycles, Index 2: 8 cycles, Read 2: 150 cycles), targeting a minimum coverage of 50,000 reads per cell.

### Data preprocessing and quality control

FASTQ alignment, filtering, and barcode counting for snRNA-seq and snATAC-seq data were performed with the ME-seq alignment pipeline (https://github.com/sai-ma-group/ME-seq-alignment-pipeline) using the human reference genome (GRCh38, GENCODE v32). We performed quality filtering based on metrics for snRNA-seq and snATAC-seq data. High-quality human brain nuclei were retained using the following criteria: (1) UMI count ranges from 500 to 20,000; (2) the number of detected genes is in the range of 300 to 8,000; (3) mitochondrial percentage < 5%; (4) number of fragments is in the range of 500 to 20,000; (5) TSS enrichment score > 1; (6) blacklist ratio < 0.1. We applied Scrublet ^74^ to measure the doublet probability and filtered potential doublets with a score > 0.15. In total, 356,721 nuclei were initially retained for further analysis.

For organoid samples, high-quality nuclei were kept based on the following criteria: 1,000 < UMI counts < 20,000, 500 < the number of detected genes < 8,000, mitochondrial percentage < 5%, 1,000 < number of fragments < 20,000, and TSS enrichment score > 3. In total, 81,838 nuclei were retained to validate molecular dynamics during early neuron development.

### Cell clustering and annotation

#### Gene expression data preprocessing and clustering

Cell clustering and annotation were performed using Scanpy (v1.10.1) ^53^. Ambient RNA was assessed using SoupX (v1.6.2) ^52^. Raw counts were normalized to the median library size and log-transformed. The top 5,000 highly variable genes (HVGs) were identified and used for principal component analysis (PCA). The first 50 principal components (PCs) were used to construct a batch-balanced nearest-neighbor graph using BBKNN ^55^. Then, Uniform Manifold Approximation and Projection (UMAP) and the Leiden algorithm were used for visualization and cell clustering, respectively. Cell classes were annotated based on the expression of canonical marker genes, such as *SLC17A7*, *GAD1*, *AQP4*, *PDGFRA*, *MBP*, *CSF1R,* and *COL1A2* for ENs, INs, Astro, OPCs, Olig, Micro_PVM, and endothelial/vascular cells, respectively.

To achieve higher-resolution annotation, ENs, INs, and fetal cells were independently reclustered using the top 3,000 HVGs and the first 30 PCs. Based on marker gene expression ^13,75^, EN subclusters were further annotated by cortical layer identity, whereas IN subclusters were classified based on their developmental origin (MGE or CGE) and neurotransmitter. Fetal ENs were annotated as radial glia (RG), intermediate progenitor cells (IPCs), newborn ENs, and immature ENs based on established marker genes. Clusters exhibiting coexpression of markers from multiple cell types were considered putative doublets and excluded from downstream analyses. In total, 340,153 high-quality nuclei were retained and classified into 29 cell subclasses.

For organoid data, cell clustering and annotation followed a similar general workflow to the primary human brain samples, with dataset-specific parameters and marker genes used to define cell identities. Cell populations were annotated based on established markers, including *SOX2, VIM, PAX6, SLC1A3,* and *NES* for RG; *FOXG1* for telencephalic identity*, SATB2, BCL11B, SLC17A6, TBR1, NEUROD2,* and *NEUROD5* for dorsal telencephalic neurons; *DLX1, DLX2, GAD1, GAD2,* and *SLC32A1* for ventral telencephalic neurons; *NR2F2, HOXB3, LMX1A,* and *LMX1B* for non-telencephalic neurons; *COL1A1, COL1A2, DCN,* and *LUM* for mesenchymal cells; and *NANOG* for pluripotent stem cells. Batch effects from donors were corrected using BBKNN ^55^. To further resolve the developmental states of organoid neurons, we performed label transfer using scVI (v1.3.0)^61^, with the primary human brain dataset as the reference.

#### ATAC and DNA methylation data processing

Preprocessing, dimensionality reduction, and clustering of ATAC and mCG data were performed using SnapATAC2 ^54^. The genome was partitioned into 5-kb bins to construct cell-by-bin matrices, using chromatin fragment counts for ATAC data and unmethylation levels for DNA methylation data. Genomic bins were selected based on total fragment counts or unmethylation levels, excluding the lowest and highest 5% of their respective distributions and those overlapping ENCODE blacklist regions ^76^. Dimensionality reduction was performed using the matrix-free spectral embedding algorithm implemented in SnapATAC2, retaining 15 spectral components, followed by batch correction using BBKNN ^55^ with three neighbors per batch. UMAP was used for visualization, and Leiden clustering was performed to identify cell clusters.

For snATAC-seq, gene activity scores were calculated by counting fragments overlapping gene bodies, and cell-type-specific accessible chromatin peaks were identified using the CallPeaks function in Signac ^56,57^, which wraps MACS2. Gene-body mCG fraction was calculated to facilitate cell-type annotation.

### Trajectory inference

For each molecular modality, trajectory inference was performed independently using Slingshot ^59^. To reconstruct EN developmental trajectories, RG, IPCs, newborn ENs, immature ENs, and layer-specific mature ENs were subsetted and reclustered. High-quality cells with an ambient RNA fraction < 0.5, number of fragments > 1000, and a TSS enrichment score > 3 were retained for cross-modal trajectory comparison. For RNA data, the top 2,000 HVGs were selected, and the effects of log-transformed total UMI counts and mitochondrial transcript percentages were regressed out. PCA was then performed, retaining the first 15 PCs. For ATAC and mCG data, up to 100,000 informative genomic bins and all eligible bins, respectively, were retained based on total fragment counts or unmethylation levels, excluding bins in the lowest and highest 5% of their respective distributions and those overlapping ENCODE blacklist regions ^76^. Spectral dimensionality reduction was performed, retaining the first 15 and 10 components for ATAC and mCG data, respectively. The resulting spectral embeddings were adjusted by regressing out log-transformed total fragment counts for ATAC data and CpG coverage for mCG data.

For oligodendroglial trajectories, a similar pipeline was applied. OPCs, immature oligodendrocytes, and mature oligodendrocytes were subsetted and reclustered. For RNA data, the top 3,000 HVGs and the first six PCs were used. For ATAC and mCG data, up to 100,000 informative genomic bins were selected for each modality, using lower and upper quantile thresholds of 0.05 and excluding ENCODE blacklist regions, followed by spectral dimensionality reduction, retaining the first six components for each modality.

For all trajectories across molecular modalities, batch correction was performed using BBKNN ^55^, followed by clustering. Slingshot was used to construct a cluster-based minimum spanning tree and fit simultaneous principal curves to infer developmental trajectories. RG and layer-specific mature EN clusters were specified as the root and terminal states for EN trajectories, respectively. Infant OPCs served as the root for oligodendroglial trajectories. Pseudotime values were assigned to individual cells based on their projections onto the principal curves and independently min–max normalized to the range of 0–1 for each lineage. The inferred trajectories were projected onto two-dimensional UMAP embeddings for visualization.

Cumulative pseudotime progression during EN development was calculated as the difference in median pseudotime between each developmental stage and RG, normalized to the total median pseudotime difference between RG and adolescent neurons. The fraction of post-infancy progression was calculated as the difference in median pseudotime between infancy and adolescence, normalized to the same total difference.

### Differential expression analysis

To identify dynamically expressed genes along each lineage, gene-wise generalized additive models (GAMs) were fitted using the negative binomial distribution in tradeSeq (v1.12.0) ^60^. The number of knots was set to four, and batch, sex, and donor source were included as covariates in the model. Dynamically expressed genes were identified using the associationTest function with an *FDR* < 0.05 and a log2 fold-change threshold of 1.

### Differential accessibility analysis

Differentially accessible regions (DARs) across development were identified using the limma-voom pipeline ^77^. For each lineage, donor-level pseudobulk profiles were generated from the cell-by-peak count matrix. Trimmed mean of M-values (TMM) normalization factors were estimated based on promoter peaks and used to calculate effective library sizes. Raw counts were then transformed to log counts per million (logCPM) using the voom function. Linear models were fitted for each peak using lmFit, and pairwise comparisons between developmental stages were performed. Empirical Bayes moderation was applied using eBayes to stabilize variance estimates. Batch, sex, and donor source were included as covariates in the design matrix. DARs were defined as regions with an *FDR* < 0.05 and an absolute log2 fold change > 1.

### Differential methylation analysis

Differentially methylated regions (DMRs) across development and cell types were identified using the DMRfind in Methylpy^78^. For developmental DMRs, pseudobulk methylation profiles were generated for each cell lineage at each developmental stage and converted to the *allc* format. DMR identification was performed independently for each lineage across developmental stages. Differentially methylated sites (DMSs) were first identified using the root mean square (RMS) test implemented in DMRfind, and sites with an *FDR* < 0.05 were considered significant. To mitigate data sparsity, neighboring cytosines within 250 bp were jointly analyzed during DMS detection. Significant DMSs within 250 bp of one another and showing consistent directions of methylation change were then merged into DMRs. DMRs were retained if they contained at least five DMSs, spanned at least 200 bp, had a mean cytosine coverage ≥ 8, and showed an absolute difference in mCG fraction (|ΔmCG|) > 0.2. For cell-type-specific DMR analysis, pseudobulk methylation profiles were similarly generated for each cell type and analyzed using the same workflow.

### Modules in dynamic features

#### DEG and DAR modules

To characterize transcriptional and chromatin accessibility dynamics across development, pseudobulk profiles of DEGs and DARs were analyzed separately for each lineage. For EN lineages, cells were grouped into eight ordered developmental bins based on age and developmental cell identity, including progenitors (RG and IPC), newborn/immature ENs, neonatal (birth), early infancy (0.2–0.5 years), early childhood (2–4 years), late childhood (5–10 years), early adolescence (12–14 years), and late adolescence (15–20 years). For the OPC lineage, cells were grouped into six ordered age-defined bins spanning neonatal, infancy, early childhood, late childhood, early adolescence, and late adolescence. For the Olig lineage, cells were grouped into seven ordered developmental bins, beginning with OPCs and followed by oligodendrocytes from neonatal, infancy, early childhood, late childhood, early adolescence, and late adolescence stages. Raw counts were normalized within each lineage using the trimmed mean of M-values (TMM) method implemented in edgeR, converted to counts per million (CPM) using the corresponding effective library sizes, and log-transformed. For each DEG and DAR, normalized values across developmental bins were modeled using a Gaussian generalized additive model (GAM) implemented in the mgcv R package. The developmental bin was modeled as a smooth term using a cubic regression spline with a maximum basis dimension of *k* = 4. Models were fitted using restricted maximum likelihood (REML), and fitted trajectories were predicted at 70 evenly spaced points across development.

#### DMR modules

For each DMR, methylated CpG counts and total CpG coverage were summarized across the lineage-specific developmental bins. Bins with a minimum CpG coverage of five were retained, and regions were required to have at least four valid developmental bins. Methylated CpG counts were modeled relative to total CpG coverage using a binomial GAM, with methylation probability modeled as a smooth function of developmental progression (*k* = 4). Models were fitted using REML, and fitted trajectories were predicted at 70 evenly spaced points across development.

To identify temporal patterns, GAM-fitted trajectories were independently min–max scaled to a range of 0–1 for each feature and subsequently grouped using k-means clustering. K-means clustering was performed with 100 random initializations and a fixed random seed of 123. The number of clusters was specified separately for each molecular modality to capture the major developmental patterns. Cluster-level trajectories were summarized by averaging the scaled fitted values of features assigned to each cluster across developmental progression.

### Relative magnitude of changes in dynamic features across development

To compare developmental remodeling across molecular modalities, DEGs, DARs, and DMRs were evaluated across ordered developmental stages within each lineage. RNA expression, chromatin accessibility, and mCG levels were independently min–max scaled to 0–1 across development for each feature. For each feature, the magnitude of change was calculated as the absolute difference in scaled signal between adjacent stages. The distributions of feature-level changes were compared across developmental transitions and molecular modalities.

### Functional enrichment analysis

Functional enrichment analysis was performed to characterize the biological processes associated with developmentally dynamic genes and regions. Gene Ontology (GO) enrichment analysis of differentially expressed genes (DEGs) was conducted using gprofiler2 (v0.2.3) ^67^. For DARs and DMRs, functional enrichment analysis was performed using rGREAT (v2.9.2) ^65^, which associates genomic regions with nearby genes and evaluates enrichment of biological annotations using the Genomic Regions Enrichment of Annotations Tool (GREAT). Gene sets containing 1,500 or more genes were excluded to improve the biological specificity of enrichment results.

### Cross-modal enrichment of developmental dynamic modules

To assess the concordance of developmental dynamics across RNA, ATAC, and mCG, DARs and DMRs were annotated to genes using ChIPseeker (v1.38.0) ^68^. Pairwise enrichment between dynamic modules across modalities was assessed at the region level using one-sided Fisher’s exact tests. Odds ratios (ORs) were calculated to quantify the strength of association, and *p*-values were adjusted for multiple testing using the Benjamini–Hochberg procedure. *FDR* < 0.05 was considered statistically significant.

### Genetic enrichment

Stratified linkage disequilibrium score regression (S-LDSC) ^62^ was applied to assess the contribution of developmentally dynamic features to the heritability of neuropsychiatric and neurological disorders. Separate annotations were constructed for different developmental patterns of DEGs, DARs, and DMRs. For gene-based annotations, genomic regions corresponding to genes within each DEG category were used to define annotation sets, whereas DARs and DMRs were directly used as region-based annotations.

S-LDSC was performed using genome-wide association study (GWAS) summary statistics for ADHD ^79^, ASD ^80^, BD ^81^, MDD ^82^, SCZ ^41^, AD ^83^, PD ^84^, and height ^85^. For each annotation, a binary SNP annotation was generated based on whether HapMap3 SNPs ^86^ overlapped the corresponding genomic regions, with SNPs in the major histocompatibility complex (MHC) region excluded from the analysis. Annotation-specific LD scores were calculated using a 1-cM window and the European-ancestry reference panel from Phase 3 of the 1000 Genomes Project ^87^. Heritability enrichment was estimated conditional on the baseline LDSC model ^62^.

The contribution of each developmental annotation to trait heritability was further quantified using the normalized S-LDSC coefficient, which represents the standardized change in per-SNP heritability associated with an annotation after conditioning on annotations included in the baseline model. Statistical significance was assessed using the *p*-value of the annotation coefficient. *p*-values were adjusted for multiple testing using the Benjamini–Hochberg procedure, and annotations with an adjusted *p* < 0.05 were considered significant. To assess the independent contributions of chromatin accessibility and DNA methylation, joint S-LDSC analyses were performed by simultaneously including the corresponding DAR and DMR annotations, together with the baseline annotations, in the same regression model.

The enrichment of risk variants of neuropsychiatric and neurological disorders across DEG modules along EN maturation was evaluated using MAGMA^63^. At the gene set level, MAGMA applied a generalized least squares regression model to test whether the genes in a gene set are more strongly associated with the GWAS signal than other genes.

### Tile–gene *cis*-regulatory association analysis

To enable directly comparable correlation testing across modalities, we adopted a TSS-centered tile-based representation rather than modality-specific peaks, as previously described ^88^. Protein-coding gene TSSs (hg38 GENCODE, n = 18,647) were flanked by ±250 kb, merged across strands, and partitioned into 500-bp tiles (n = 4,229,294). For each chromatin modality, tiles with non-zero signal across pseudobulks were retained, ENCODE hg38 blacklist regions were excluded, GC content was annotated using the chromVAR package, and tiles with undefined GC content were discarded.

#### Multimodal pseudobulk aggregation

To address single-cell methylation data sparsity and ensure matched dimensions, all three modalities were aggregated into matched pseudobulks. Within each sample x cell subtype stratum, cells were randomly partitioned into non-overlapping pseudobulks of 100 cells. Strata containing fewer than 100 cells were collapsed into a single pseudobulk.

#### Background-matched correlation testing

Candidate tile–gene associations were tested against covariate-matched backgrounds as previously described ^20,29^. A background pool of 10^5^ tile–gene pairs was randomly sampled. Each candidate pair was matched to the 100 most similar background pairs based on tile GC content, mean tile signal, and mean gene expression. Spearman correlation was tested against the mean and standard deviation of matched background correlations using a *Z*-test. Tiles mapping to multiple genes were retained.

For mCG–RNA, *p*-values were derived from a two-tailed Z-test to account for both positive and negative associations, whereas ATAC–RNA used one-tailed tests. Uncovered mCG values were imputed with the mean methylation rate across covered pseudobulks prior to correlation testing.

The pipeline was executed on the full dataset and each cell-type subset. Significant links were defined by *p* < 0.01 and cell-type-specific Spearman correlation cutoffs (**Fig. S2A** and **S2B**). Endothelial/vascular and RG subsets were excluded from downstream analyses due to limited pseudobulk numbers and unstable correlations.

DORC definition was restricted to tile–gene links within ±50 kb of TSSs to minimize false-positive long-range associations. For each modality, significant links from the full dataset and cell-type-stratified analyses were pooled and deduplicated by tile–gene pair. DORC genes were identified based on the number of significant tiles associated with each gene, which was set at the inflection point of the ranked link-count distribution.

#### PhastCons conservation enrichment of linked tiles

PhastCons 100-way vertebrate conservation scores were retrieved using the GenomicScores R package and averaged across each 500-bp tile. Significant tile–gene associations within ±50 kb of TSSs defined the linked-tile sets for each modality, with tiles absent from all significant links serving as background. Conservation distributions were compared between each linked group and the unlinked background using one-sided Wilcoxon rank-sum tests with Benjamini-Hochberg correction.

#### Functional enrichment of DORC genes

ATAC- and mCG-derived DORC gene sets were combined by union. GO Biological Process 2025 enrichment was performed using the enrichR package, with all protein-coding genes detected in the RNA matrix as background. The top 30 GO terms ranked by adjusted *p*-value were grouped into functional modules by manual assignment to high-level parent categories of the GO Biological Process hierarchy. GO terms with a gene overlap coefficient |A ∩ B| / min(|A|, |B|) ≥ 0.10 were connected in a network visualized using igraph and ggraph packages. The top 30 GO Biological Process terms were scored using the AddModuleScore function in Seurat. Scores were averaged by cell type and z-score standardized across cell types within each term.

#### Enrichment of DORC genes in super-enhancer-associated gene sets

The unified DORC gene set was tested for enrichment in two independent SE annotations ^15,22^. Hypergeometric tests were performed for each tissue or brain cell type using detected protein-coding genes as background. *p*-values were adjusted across tissues or cell types using the Benjamini–Hochberg correction.

#### Multimodal heatmap representation of tile–gene links

Significant links within ±50 kb of TSSs shared by ATAC–RNA and mCG–RNA were used for visualization. Per-tile TF–IDF-normalized accessibility, per-tile mCG rate, and per-gene log-normalized expression were extracted at the pseudobulk level for each link. The accessibility and expression matrices were row-wise z-score standardized, and mCG rate matrices were retained on the original 0–1 scale. Links were clustered into 10 groups by k-means on the z-scored accessibility matrix. Three matrices for the same set of links were rendered side-by-side in the cluster-resolved row order. Pseudobulk columns were ordered by binary sorting of cluster centroids and annotated with cell type, cell subtype, developmental stage, and donor age.

For each k-means cluster, the 200 genes most frequently linked were tested for GO Biological Process enrichment using the topGO R package, with all genes appearing in any cluster of the link set used as the gene universe. Top terms were ranked by enrichment *p*-value.

Link clusters were annotated using published cell-type-specific ChromHMM segmentations from the human brain for neuronal, glial, and fetal categories ^23^. For the integrated link set, clusters 1–6 were assigned to glia, clusters 7–8 to neurons, and clusters 9–10 were excluded from this analysis. For the EN-specific link set, cluster 3 was assigned to the fetal category and the remaining clusters were assigned to the neuronal category. Each tile was then assigned the ChromHMM state with the largest base-pair overlap using the GenomicRanges R package. State enrichment relative to all annotated links was assessed by Fisher’s exact test. Effect sizes were expressed as log_2_(observed proportion/background proportion), and *p*-values were adjusted using the Benjamini-Hochberg correction.

To assess the spatial interactions of linked tiles, NeuN+ neuronal PLAC-seq chromatin loops were obtained from Nott et al. ^15^. Significant linked tiles were intersected with PLAC-seq loop anchors based on genomic coordinates.

### Developmental clock modeling

#### Bootstrap pseudocell construction

Single cells were aggregated into bootstrap pseudocells stratified by donor and cell type, employing the BootstrapCell strategy introduced by Buckley et al. ^89^. Strata containing at least 1,500 cells were sampled without replacement. Smaller strata were sampled with replacement to preserve the target pseudocell size and number. Each pseudocell comprised 50 cells, with 30 pseudocells generated per stratum. These parameters were selected by benchmarking inter-fold stability (1 / coefficient of variation of fold-wise MAE) and donor-level Pearson R^2^ of predicted vs. true log-age across modalities.

#### Modality-specific input matrices

For the RNA input matrix, the top 5,000 highly variable protein-coding genes were selected using the Seurat R package, and pseudocell expression was computed as the mean of log-normalized values across the constituent cells. For the ATAC input matrix, the input feature set was 500-bp tiles significantly associated with any of these genes within ±250 kb. Pseudocell accessibility was averaged from TF–IDF-normalized values. For the mCG input matrix, the input feature set was 500-bp tiles significantly associated with any of these genes within ±250 kb, filtered by cohort-wide cytosine coverage > 0.1%. Methylation rates were averaged across constituent cells with non-zero coverage. Missing mCG values were hierarchically imputed using per-tile means at the donor–cell-type, developmental-stage–cell-type, cell-type, and global levels, with imputation performed separately within cross-validation folds to avoid leakage. Cell-type-specific models used top 1,000 highly variable protein-coding genes for RNA and their significantly associated tiles derived from the cell type-stratified analysis for ATAC and mCG, with cell-type-specific mCG coverage filtering.

#### Model training and evaluation

For each modality, LASSO regression was performed using the glmnet R package to predict log-age based on pseudocell features. Models were evaluated using five-fold cross-validation. For each held-out group, the model was trained on remaining groups, with λ selected by inner cross-validation minimizing MAE. Predicted log-ages were aggregated using the median within each donor and validation fold, and performance was assessed by MAE and Pearson R^2^.

#### Cell-type-specific developmental rates

Predicted log-ages were summarized by median for each cell type and age. For each modality and cell type, a smoothing spline was fit to the predicted log-age as a function of true log-age. To quantify the cell-type-specific developmental rates, the fitted spline was evaluated at 100 equally spaced points across the observed age range. Developmental rate was defined as the mean slope between adjacent points. Rates were ranked across cell types within each modality.

#### Functional enrichment of features selected by developmental clock models

Genes with non-zero coefficients were extracted from pooled and cell-type-specific models. For ATAC and mCG models, tiles were mapped to linked genes via the significant tile–gene associations. Functional enrichment was performed using the enrichR package against a background of all detected protein-coding genes.

### Modality temporal order along EN pseudotime

After excluding fetal cells, the EN L2/3 IT lineage pseudotime trajectory was discretized into 20 equally populated bins by quantiles of the RNA-derived pseudotime. Cells within each bin were aggregated using the corresponding modality-specific preprocessing procedures. Predicted log-ages were generated for each pseudocell by the corresponding EN-specific developmental clock model, yielding modality-specific developmental trajectories. Modality differences across matched pseudotime bins were assessed using paired Wilcoxon signed-rank tests with Benjamini–Hochberg correction.

### Gene regulatory networks

#### TF motif activity inference

TF motif activities were inferred independently for ATAC and mCG modalities using the chromVAR framework, restricted to EN L2/3 IT lineage cells. For ATAC, GC bias of peaks was estimated, and GC-content- and accessibility-matched background peak sets were generated. TF motif occurrences were identified using the human PFM collection from chromVARmotifs with the motifmatchr R package. Motif deviation Z-scores were computed using the chromVAR R package. For mCG, motif matching and background peak selection were performed on the corresponding ATAC peak coordinates. Methylation-based motif Z-scores were then computed using a custom adaptation of the chromVAR algorithm, in which peak-level expectations were defined as coverage-weighted regional methylation rates, and per-sample scaling factors were defined as per-cell mean methylation rates. Otherwise, deviation, mean-and-SD-based Z-score normalization, and background subtraction followed the original chromVAR formulation. The resulting Z-score matrices were column-wise scaled for both modalities.

#### DORC score and RNA expression smoothing

EN-specific DORCs were used as the input gene set. DORC scores were computed by summarizing the constituent linked tiles per gene. ATAC scores were defined as the mean tile signal across linked tiles, while mCG scores were defined as the coverage-weighted regional methylation rate across linked tiles. To denoise the single-cell scores, a 30-nearest-neighbor graph was constructed in the cell PCA space. For the ATAC DORC score and RNA expression matrix, smoothing was performed by averaging across each cell and its 30 neighbors. For the mCG DORC score matrix, smoothing was instead performed only over neighbors with non-zero coverage at each tile, followed by per-tile mean imputation of any remaining uncovered entries.

#### FigR regulatory inference

TF–DORC regulatory interactions were inferred independently for the ATAC and mCG modalities using FigR ^29^, which combines TF motif enrichment within the cis-regulatory tiles linked to a given DORC and its 30 nearest DORCs in scaled DORC space, with the Spearman correlation between the smoothed DORC score and the smoothed TF expression across cells. The two complementary statistics are combined into a signed Regulation Score, where the sign is determined by the correlation and the magnitude by the combined enrichment and correlation *p*-values. The merged network was rendered with igraph and ggraph R packages.

