## Supplementary figures for "Tri-Modal Single-Cell Atlas of Developing Human Brain"

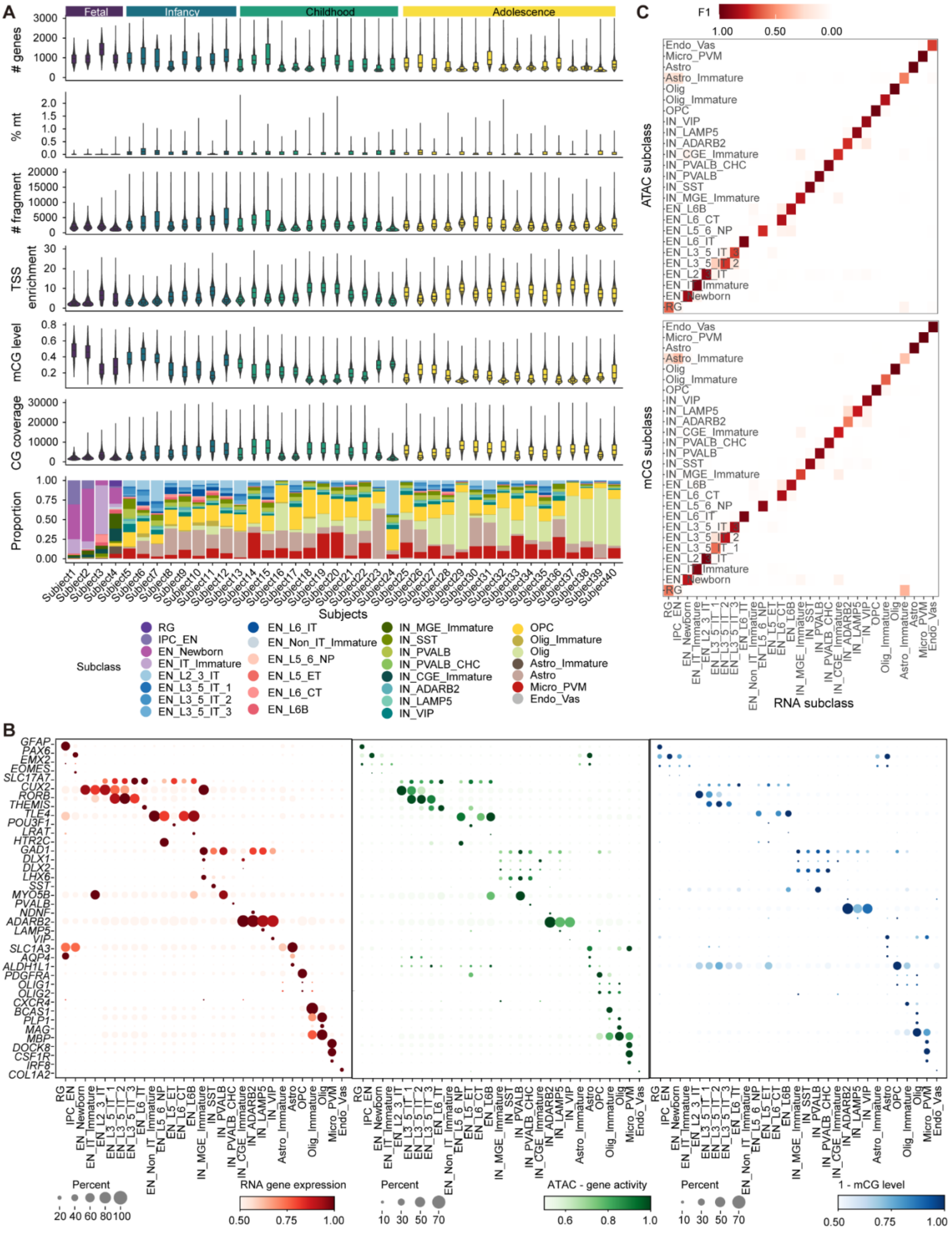
A
Fetal
Infancy
Childhood
Adolescence
### genes
% mt
### fragment
TSS enrichment
mCG level
CG coverage
Proportion
Subjects
Subclass
RG
IPC_EN
EN_Newborn
EN_IT_Immature
EN_L2_3_IT
EN_L3_5_IT_1
EN_L3_5_IT_2
EN_L3_5_IT_3
EN_L6_IT
EN_Non_IT_Immature
EN_L5_6_NP
EN_L5_ET
EN_L6_CT
EN_L6B
IN_MGE_Immature
IN_SST
IN_PVALB
IN_PVALB_CHC
IN_CGE_Immature
IN_ADARB2
IN_LAMP5
IN_VIP
OPC
Olig_Immature
Olig
Astro_Immature
Astro
Micro_PVM
Endo_Vas
B
RNA gene expression
ATAC - gene activity
1 - mCG level
Percent
C
F1
ATAC subclass
mCG subclass
RNA subclass

**Figure S1. Single-nucleus multi-omic atlas of human brain development, related to Figure 1. (A)** Quality-control metrics and cell-type composition across 40 human brain samples spanning fetal development, infancy, childhood, and adolescence. Violin plots show the distributions of detected gene counts and mitochondrial transcript fractions for snRNA-seq, fragment counts and transcription start site (TSS) enrichment scores for snATAC data, and global mCG fraction and CG coverage for sn-mCG data. The stacked bar plot shows the relative proportions of annotated cell subclasses in each sample. Samples are ordered by developmental age. **(B)** Marker-gene expression (left), ATAC-derived gene activity (middle), and unmethylated CG fraction (1 – mCG fraction; right) across annotated subclasses. Dot size represents the percentage of nuclei with detected signal, and color represents the scaled average signal for each modality. **(C)** Concordance of cell-subclass annotations across modalities. Heatmaps show pairwise F1 scores between RNA- and ATAC-based subclass annotations (top) and RNA- and mCG-based subclass annotations (bottom).

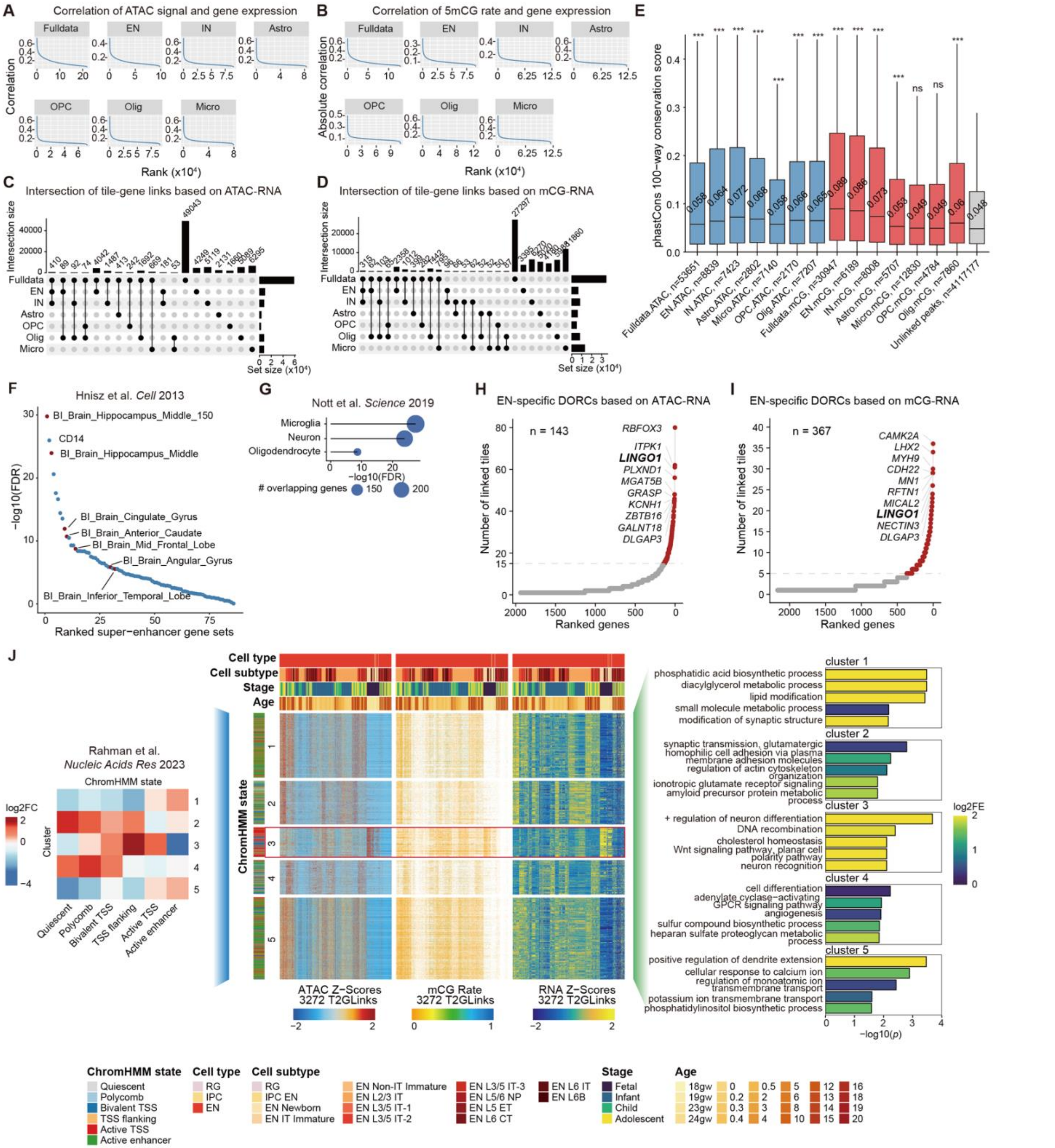

**Figure S2. Validation and cell-type-stratified analyses of tile–gene cis-regulatory links, related to Figure 2. (A)** Ranked Spearman correlation coefficients (ρ) for ATAC–RNA links ($p < 0.01$, $\rho > 0$) within ±50 kb of the TSS, across cell types and the full dataset. A cutoff of $\rho \geq 0.2$ was selected at the curve inflection point for DORC definition. **(B)** As in (A), for mCG–RNA links ($p < 0.01$; both positive and negative correlations retained). A cutoff of $|\rho| \geq 0.15$ was selected at the curve inflection point for DORC definition. **(C)** UpSet plot of significant ATAC–RNA tile–gene links across cell types and the full dataset. Horizontal bars indicate the total number of links per group. Vertical bars indicate the size of each intersection. **(D)** As in (C), for significant mCG–RNA tile–gene links. **(E)** PhastCons 100-way vertebrate conservation scores for linked versus unlinked tiles. Linked tiles were derived from significant ATAC–RNA or mCG–RNA associations (±50 kb), across the full dataset and cell types. Tiles absent from any significant link served as the background. Boxplots show the group distribution, with median values annotated. One-sided Wilcoxon rank-sum tests were used, with Benjamini–Hochberg correction. *** adjusted $p < 0.001$. **(F)** Enrichment of the unified DORC genes in tissue-specific super-enhancer-associated gene sets. Each point represents one tissue. Y-axis: $-\log_{10}(FDR)$ from a one-sided hypergeometric test using all detected protein-coding genes as background. Benjamini–Hochberg correction was applied across tissues. Brain-related tissues are highlighted in red. **(G)** Enrichment of the unified DORC genes in brain cell-type-specific gene sets distally linked to super-enhancers. Dot size indicates the number of overlapping genes. X-axis: $-\log_{10}(FDR)$ from a one-sided hypergeometric test. Benjamini–Hochberg correction was applied across cell types. **(H)** Ranked distribution of significantly linked tiles per gene for EN-specific ATAC–RNA associations (±50 kb). Genes with ≥ 15 significant tiles were designated as EN-specific ATAC–RNA DORCs (n = 143). Top DORC genes are labeled. **(I)** As in (H), for EN-specific mCG–RNA associations. Genes with ≥ 5 significant tiles were designated as EN-specific mCG–RNA DORCs (n = 367). **(J)** As in Figure 2H, EN-specific ATAC–RNA tile–gene links (n = 3,272) were clustered into 5 groups. Clusters were annotated using fetal (cluster 3) or neuronal (remaining clusters) ChromHMM segmentations. Column annotations indicate cell subtype, developmental stage, and donor age.

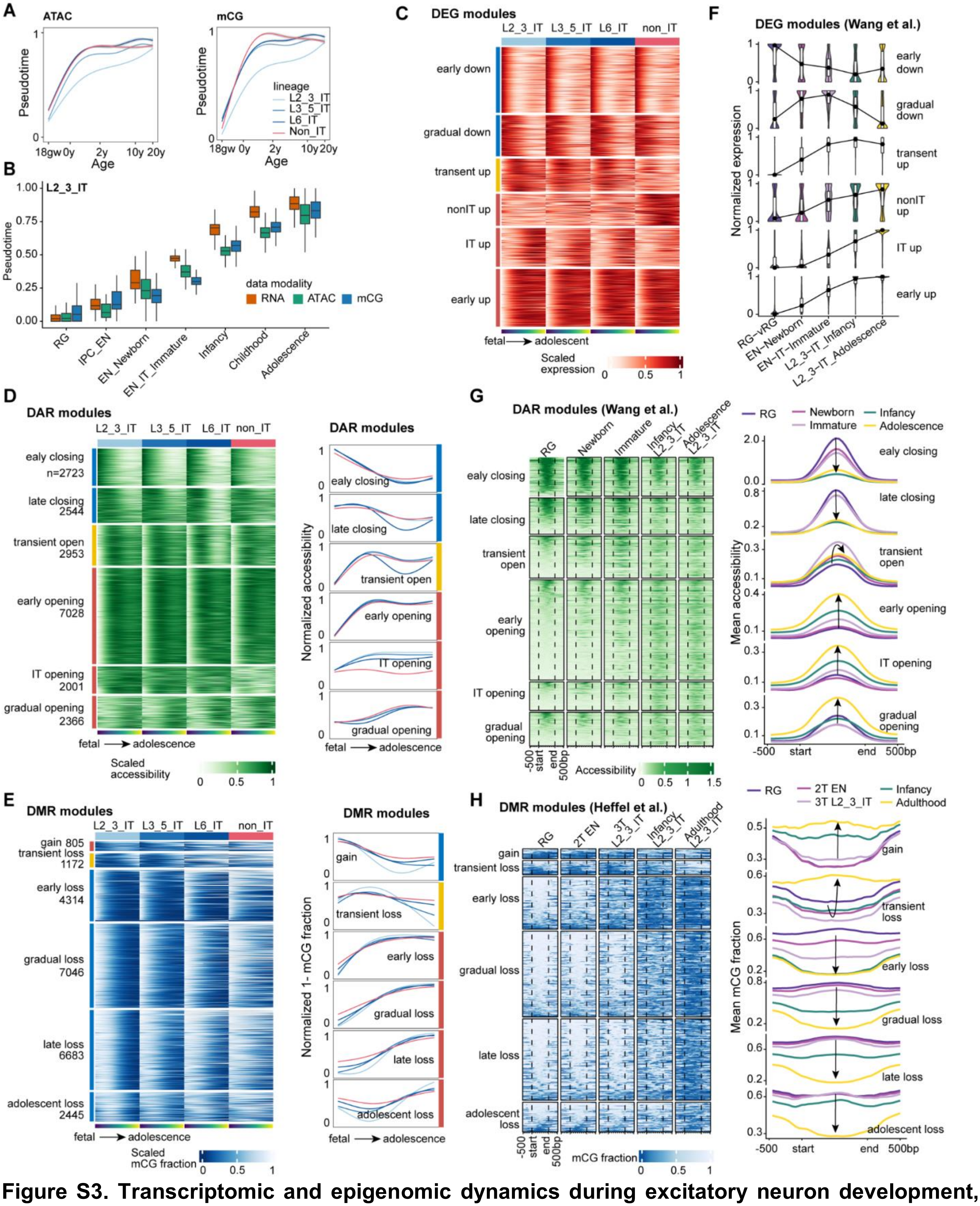


**Figure S3. Transcriptomic and epigenomic dynamics during excitatory neuron development, related to Figure 3. (A)** Relationships between ATAC- (left) and mCG-derived (right) pseudotime and age across EN lineages. Curves were fitted using generalized additive models (GAMs). **(B)** Distributions of RNA-, ATAC-, and mCG-derived pseudotime across developmental cell states and stages along the L2/3 IT lineage. **(C)** Differentially expressed genes (DEGs) across development from fetal to adolescent stages in L2/3 IT, L3/5 IT, L6 IT, and non-IT lineages. Heatmaps show min–max scaled expression for each DEG. **(D)** Developmentally dynamic differentially accessible regions (DARs), with mean normalized accessibility for each module shown on the right. Numbers indicate the number of DARs in each module. **(E)** Developmentally differentially methylated regions (DMRs), with mean normalized unmethylated CG (1 – mCG fraction) profiles for each module shown on the right. Numbers indicate the number of DMRs in each module. **(F–H)** Validation of developmental molecular dynamics using independent human brain datasets. **(F)** Normalized expression of genes within each DEG module across EN development in the Wang et al. dataset [1]. **(G)** Chromatin accessibility at DARs within each module and their 500-bp flanking regions across EN L2/3 IT development in the Wang et al. dataset [1], with mean profiles shown on the right. Arrows indicate progression from radial glia to adolescent EN L2/3 IT. **(H)** mCG levels at DMRs within each module and their 500-bp flanking regions across EN development in the Heffel et al. dataset [2], with mean profiles shown on the right. Arrows indicate progression from radial glia to adult L2/3 IT neurons. 2T, second trimester; 3T, third trimester.

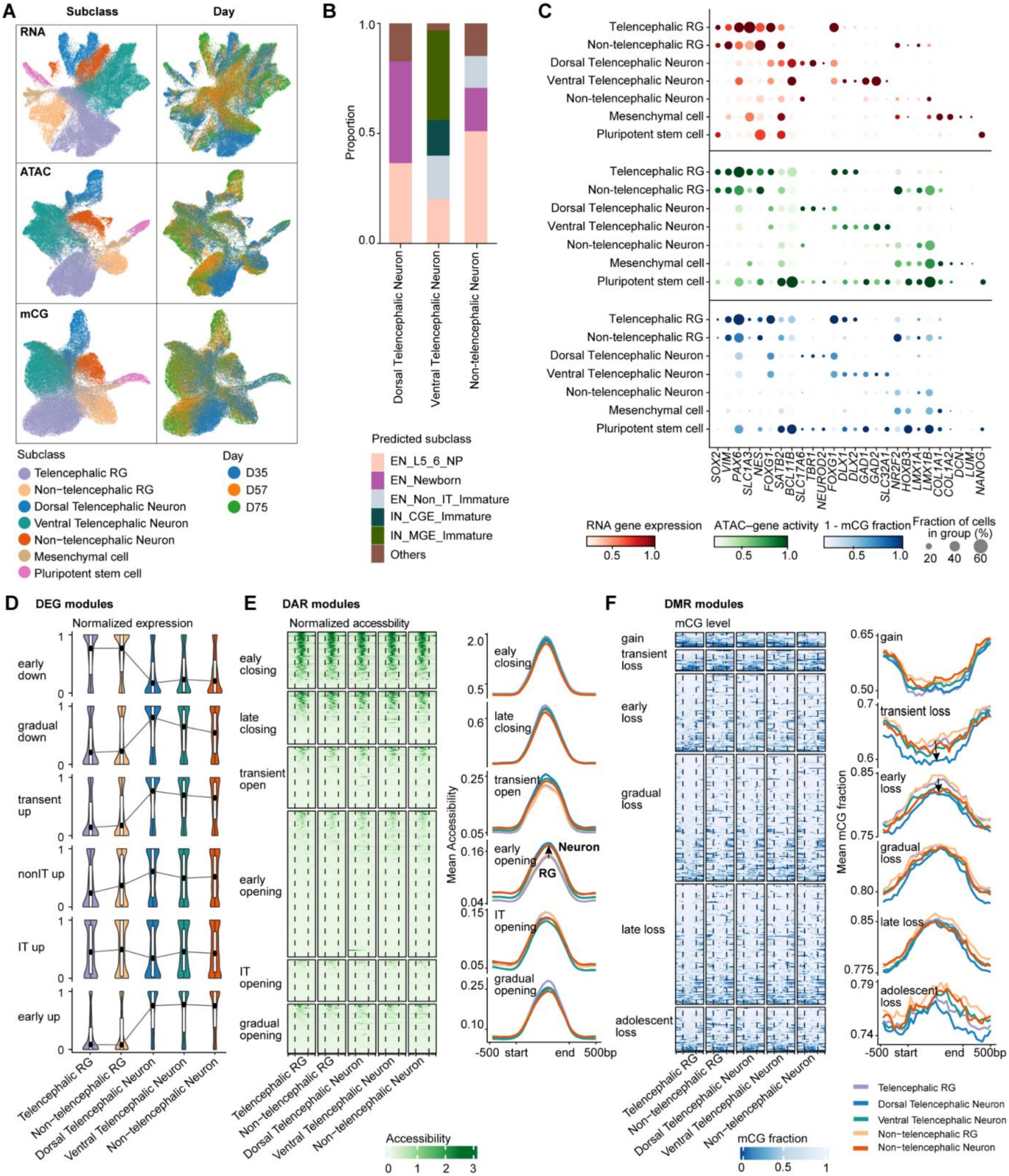


**Figure S4. Organoid ME-seq profiling captured early molecular dynamics during neuronal differentiation, related to Figure 3. (A)** UMAP visualization of organoid cells based on RNA expression, chromatin accessibility (ATAC), and DNA methylation (mCG), colored by cell subclass or developmental day (D35, D57, and D75). **(B)** Predicted neuronal subclass composition across dorsal telencephalic, ventral telencephalic, and non-telencephalic neurons. **(C)** Marker-gene expression, ATAC-derived gene activity, and unmethylated CG fraction (1 − mCG fraction) across annotated cell subclasses. Dot size indicates the fraction of cells in each group. **(D–F)** Organoid profiles of developmental DEG, DAR, and DMR modules identified in the primary human brain. **(D)** Normalized expression of genes within each DEG module across RG and neuronal populations. **(E)** Chromatin accessibility across DARs and their 500-bp flanking regions for each module, with mean profiles shown on the right. Arrows indicate progression from radial glia to neurons. **(F)** mCG fraction across DMRs and their 500-bp flanking regions for each module, with mean profiles shown on the right.

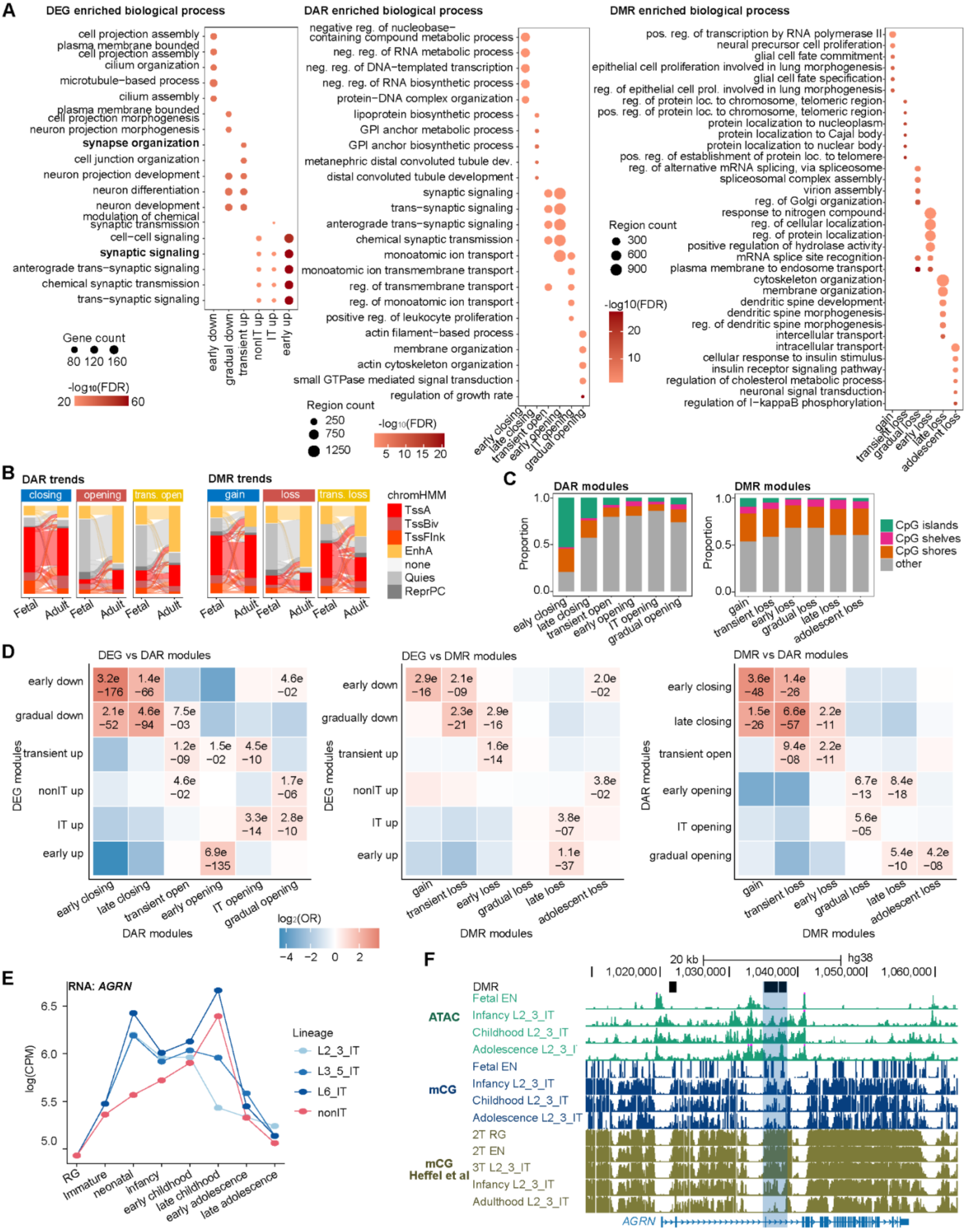

A
DEG enriched biological process
cell projection assembly
plasma membrane bounded cell projection assembly
cilium organization
microtubule−based process
cilium assembly
plasma membrane bounded cell projection morphogenesis
neuron projection morphogenesis
synapse organization
cell junction organization
neuron projection development
neuron differentiation
neuron development
modulation of chemical synaptic transmission
cell−cell signaling
synaptic signaling
anterograde trans−synaptic signaling
chemical synaptic transmission
trans−synaptic signaling
early down
gradual down
transient up
nonIT up
IT up
early up
Gene count
80 120 160
-log10(FDR)
20
60
DAR enriched biological process
negative reg. of nucleobase−containing compound metabolic process
neg. reg. of RNA metabolic process
neg. reg. of DNA−templated transcription
neg. reg. of RNA biosynthetic process
protein−DNA complex organization
lipoprotein biosynthetic process
GPI anchor metabolic process
GPI anchor biosynthetic process
metanephric distal convoluted tubule dev.
distal convoluted tubule development
synaptic signaling
trans−synaptic signaling
anterograde trans−synaptic signaling
chemical synaptic transmission
monoatomic ion transport
monoatomic ion transmembrane transport
reg. of transmembrane transport
reg. of monoatomic ion transport
positive reg. of leukocyte proliferation
actin filament−based process
membrane organization
actin cytoskeleton organization
small GTPase mediated signal transduction
regulation of growth rate
Region count
250
750
1250
-log10(FDR)
5 10 15 20
early closing
late closing
transient open
early opening
IT opening
gradual opening
DMR enriched biological process
pos. reg. of transcription by RNA polymerase II
neural precursor cell proliferation
glial cell fate commitment
epithelial cell proliferation involved in lung morphogenesis
glial cell fate specification
reg. of epithelial cell prol. involved in lung morphogenesis
reg. of protein loc. to chromosome, telomeric region
pos. reg. of protein loc. to chromosome, telomeric region
protein localization to nucleoplasm
protein localization to Cajal body
protein localization to nuclear body
pos. reg. of establishment of protein loc. to telomere
reg. of alternative mRNA splicing, via spliceosome
spliceosomal complex assembly
virion assembly
reg. of Golgi organization
response to nitrogen compound
reg. of cellular localization
reg. of protein localization
positive regulation of hydrolase activity
mRNA splice site recognition
plasma membrane to endosome transport
cytoskeleton organization
membrane organization
dendritic spine development
dendritic spine morphogenesis
reg. of dendritic spine morphogenesis
intercellular transport
intracellular transport
cellular response to insulin stimulus
insulin receptor signaling pathway
regulation of cholesterol metabolic process
neuronal signal transduction
regulation of I−kappaB phosphorylation
Region count
300
600
900
-log10(FDR)
20
10
gain
transient loss
gradual loss
early loss
late loss
adolescent loss
B
DAR trends
closing
opening
trans. open
DMR trends
gain
loss
trans. loss
Fetal
Adult
chromHMM
TssA
TssBiv
TssFlnk
EnhA
none
Quies
ReprPC
C
DAR modules
DMR modules
Proportion
1.0
0.5
0.0
ealy closing
late closing
transient open
early opening
IT opening
gradual opening
gain
transient loss
early loss
gradual loss
late loss
adolescent loss
CpG islands
CpG shelves
CpG shores
other
D
DEG vs DAR modules
DEG vs DMR modules
DMR vs DAR modules
early down
gradual down
gradually down
transient up
nonIT up
IT up
early up
DEG modules
DAR modules
DMR modules
early closing
late closing
transient open
early opening
IT opening
gradual opening
3.2e −176
1.4e −66
4.6e −02
2.1e −52
4.6e −94
7.5e −03
1.2e −09
1.5e −02
4.5e −10
4.6e −02
1.7e −06
3.3e −14
2.8e −10
6.9e −135
2.9e −16
2.1e −09
2.0e −02
2.3e −21
2.9e −16
1.6e −14
3.8e −02
3.8e −07
1.1e −37
3.6e −48
1.4e −26
1.5e −26
6.6e −57
2.2e −11
9.4e −08
2.2e −11
6.7e −13
8.4e −18
5.6e −05
5.4e −10
4.2e −08
log2(OR)
−4 −2 0 2
E
RNA: AGRN
log(CPM)
6.5
6.0
5.5
5.0
RG
Immature
neonatal
infancy
early childhood
late childhood
early adolescence
late adolescence
Lineage
L2_3_IT
L3_5_IT
L6_IT
nonIT
F
20 kb
hg38
1,020,000
1,030,000
1,040,000
1,050,000
1,060,000
DMR
ATAC
Fetal EN
Infancy L2_3_IT
Childhood L2_3_IT
Adolescence L2_3_IT
mCG
mCG Heffel et al
2T RG
2T EN
3T L2_3_IT
Adulthood L2_3_IT
AGRN

**Figure S5. Functional enrichment and cross-modal correspondence of dynamic regulatory modules during excitatory neuron development, related to Figure 3. (A)** Gene Ontology (GO) biological process enrichment of DEG modules (left), DAR modules (middle), and DMR modules (right). Reg.: regulation, pos.: positive, neg.: negative, dev.: development, prol.: proliferation. **(B)** Chromatin-state transitions from fetal brain to adult neurons across DAR (left) and DMR (right) patterns. Trans, transient; EnhA, active enhancer; Quies, quiescent; ReprPC, Polycomb-repressed; TssA, active transcription start site; TssBiv, bivalent/poised TSS; TssFlnk, TSS-flanking region. **(C)** CpG-context composition of developmental DAR and DMR modules. **(D)** Pairwise enrichment between dynamic modules across molecular modalities, including DEG versus DAR (left), DEG versus DMR (middle), and DMR versus DAR (right). One-sided Fisher's exact test was applied to determine the odds ratio (OR) and significance. **(E)** Pseudobulk expression of *AGRN* across EN maturation. **(F)** Genomic tracks showing ATAC and mCG profiles of the *AGRN* locus across L2/3 IT development. The blue block highlights DMRs exhibiting continued methylation remodeling.

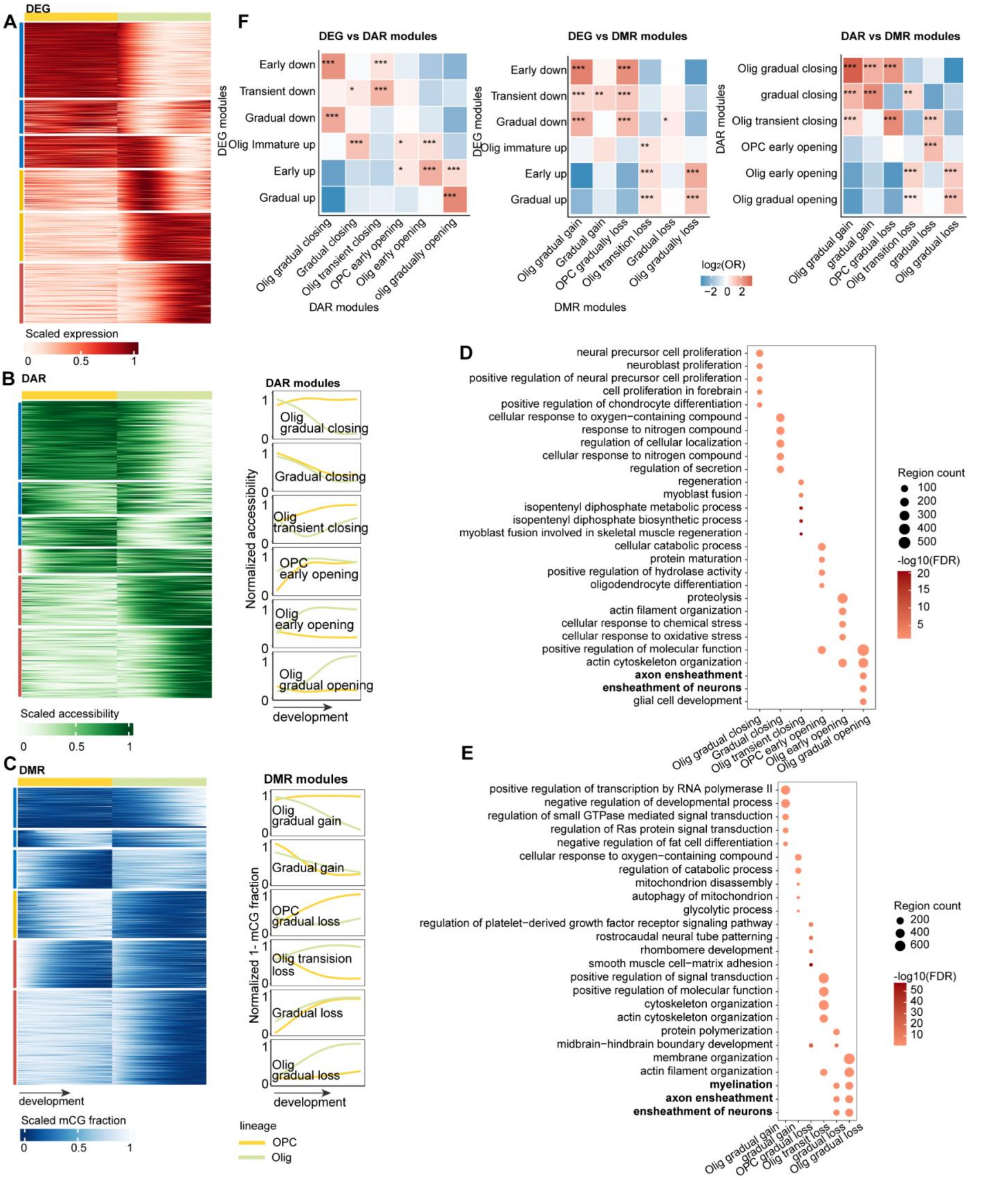


**Figure S6. Transcriptomic and epigenomic dynamics across oligodendroglial maturation, related to Figure 4. (A)** Developmental expression patterns of DEGs across oligodendroglial maturation. Heatmaps show min–max-scaled expression. **(B)** Heatmap of scaled accessibility across six developmental DAR modules, with mean normalized accessibility profiles shown on the right. **(C)**

Heatmap of scaled methylation levels across six developmental DMR modules. Line plots show mean normalized unmethylated CG fractions (1 – mCG%) along oligodendroglial maturation. **(D)** Gene Ontology (GO) biological process enrichment of DAR modules. **(E)** GO biological process enrichment of DMR modules. **(F)** Pairwise enrichment between dynamic modules across molecular modalities, including DEG versus DAR (left), DEG versus DMR (middle), and DMR versus DAR (right). One-sided Fisher's exact test was applied to determine the odds ratio (OR) and significance. * *FDR* < 0.05, ** *FDR* < 0.01, *** *FDR* < 0.001.

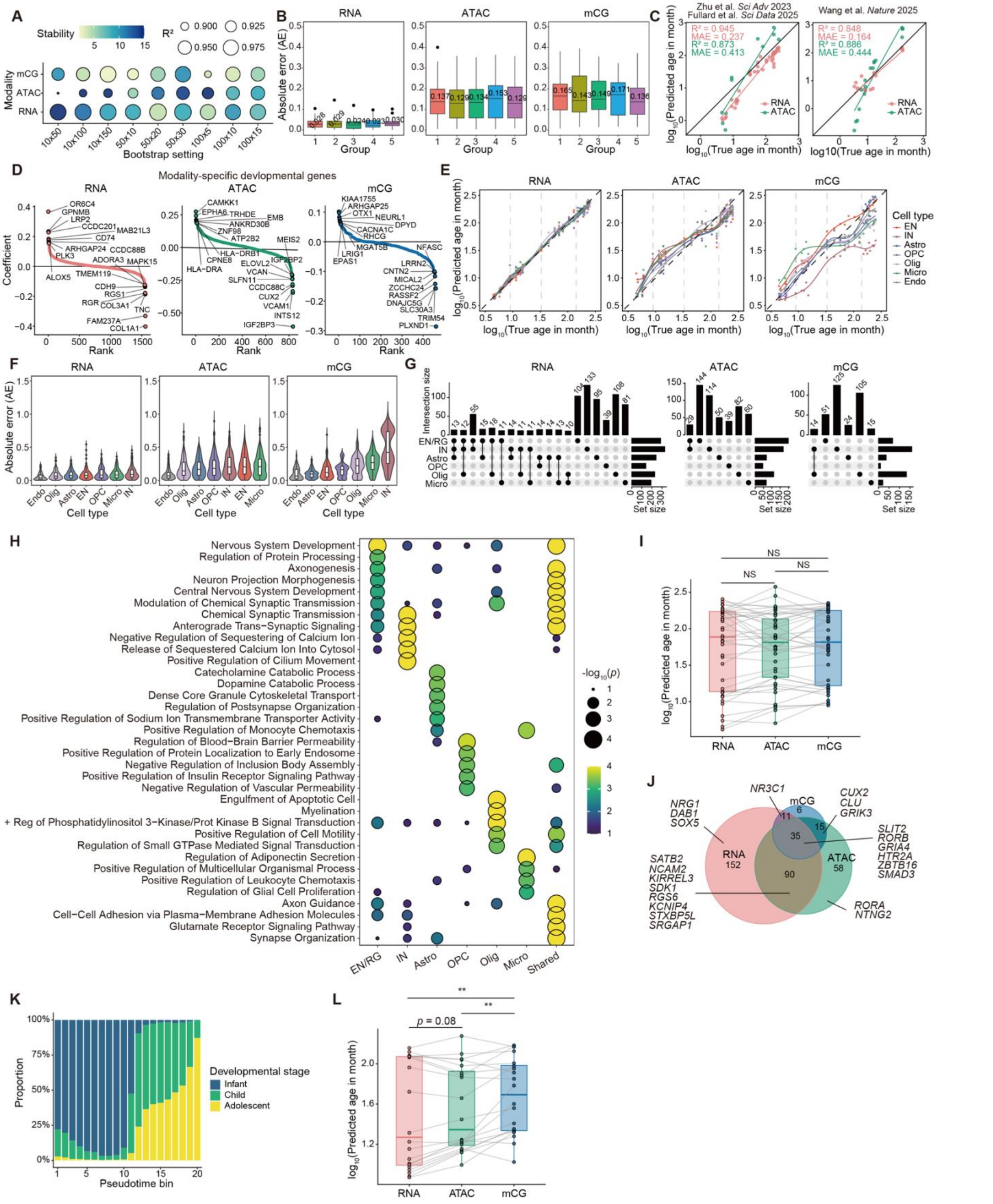


**Figure S7. Benchmark, validation, and cell-type-stratified analyses of developmental clock models, related to Figure 5. (A)** Bootstrap parameter benchmark across nine combinations of pseudocell size (number of cells per pseudocell) and n (number of pseudocells per stratum). Dot size indicates donor-level $R^2$. Color indicates inter-fold stability (1 / CV of fold-wise MAE). The selected setting (size = 50, n = 30) jointly optimized both metrics across modalities. **(B)** Per-fold absolute error (AE) distributions for cell-type-pooled RNA, ATAC, and mCG models. Each boxplot represents one held-out fold. Median AE values are annotated. **(C)** External validation of the cell-type-pooled RNA and ATAC developmental clocks on independent datasets. Each point represents one donor. $R^2$ and MAE are shown. **(D)** Ranked non-zero coefficient genes for the cell-type-pooled RNA, ATAC, and mCG models. The y-axis shows the LASSO coefficient. Top positive and negative coefficient genes are labeled. **(E)** Cell-type-resolved predicted versus true log-age from the cell-type-pooled developmental clock. Each point represents the median predicted log-age across pseudocells for a given donor and cell type. Cell-type-specific smoothing splines were used to derive developmental rates (Figure 5D). Vertical dashed lines indicate developmental stage boundaries. **(F)** Cell-type-specific AE distributions for cell-type-pooled RNA, ATAC, and mCG models. Violin/boxplots show the distributions of pseudocell-level AE. **(G)** UpSet plots of non-zero coefficient genes across cell-type-specific models, shown separately for RNA, ATAC, and mCG. Horizontal bars indicate the total number of genes per cell type. Vertical bars indicate intersection sizes. **(H)** GO biological process enrichment of genes selected by cell-type-specific developmental clock models, pooled across modalities within each cell type. Genes unique to one cell type were assigned to that cell type, while genes found in multiple cell types were grouped as shared. **(I)** Paired comparison of donor-level predicted log-ages across modalities using the EN/RG-specific developmental clock shown in Figure 5F. Lines connect matched donors across modalities. Paired Wilcoxon signed-rank tests with Benjamini–Hochberg correction. NS: not significant. **(J)** Venn diagram of non-zero coefficient genes across modalities for the EN/RG-specific model. Representative modality-specific and shared genes are labeled. **(K)** Developmental stage composition of the 20 pseudotime bins defined for the analysis shown in Figure 5I. **(L)** Paired comparison of predicted log-ages across modalities for all 20 pseudotime bins. Lines connect matched bins. Paired Wilcoxon signed-rank tests with Benjamini–Hochberg correction. **adjusted $p < 0.01$.

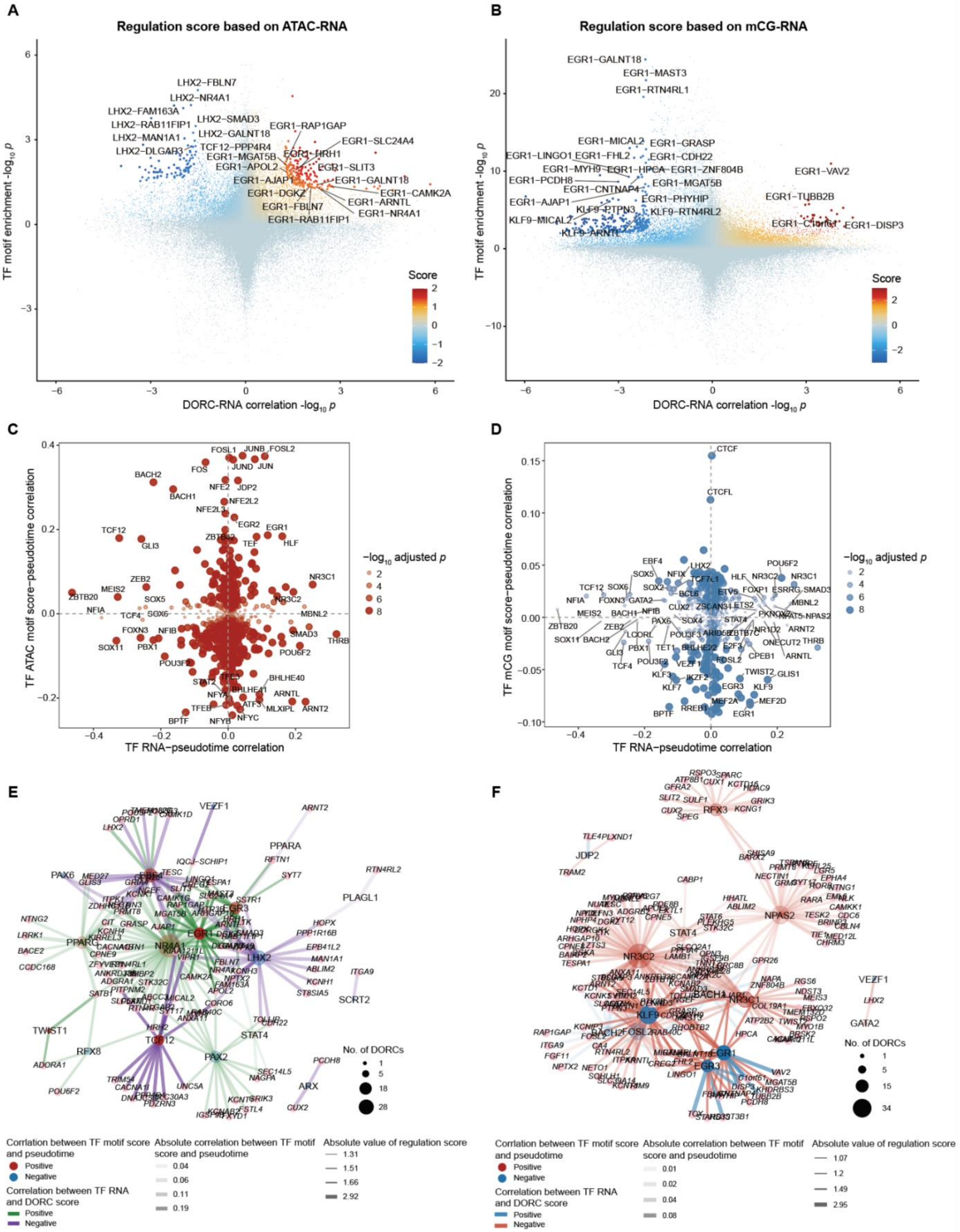
A
Regulation score based on ATAC-RNA
TF motif enrichment -log10 p
DORC-RNA correlation -log10 p
Score
B
Regulation score based on mCG-RNA
TF motif enrichment -log10 p
DORC-RNA correlation -log10 p
Score
C
TF ATAC motif score–pseudotime correlation
TF RNA–pseudotime correlation
−log10 adjusted p
D
TF mCG motif score–pseudotime correlation
TF RNA–pseudotime correlation
−log10 adjusted p
E
No. of DORCs
Corrlation between TF motif score and pseudotime
Positive
Negative
Correlation between TF RNA and DORC score
Positive
Negative
Absolute correlation between TF motif score and pseudotime
0.04
0.06
0.11
0.19
Absolute value of regulation score
1.31
1.51
1.66
2.92
F
No. of DORCs
Corrlation between TF motif score and pseudotime
Positive
Negative
Correlation between TF RNA and DORC score
Positive
Negative
Absolute correlation between TF motif score and pseudotime
0.01
0.02
0.04
0.08
Absolute value of regulation score
1.07
1.2
1.49
2.95

**Figure S8. TF motif–pseudotime correlations and modality-specific regulatory network details, related to Figure 6. (A)** Global FigR regulation profile based on ATAC–RNA within the EN L2/3 IT lineage. Each point represents a TF–DORC pair. X-axis: signed correlation significance between TF RNA expression and DORC score. Y-axis: signed significance of TF motif enrichment within linked tiles. Color indicates regulation score. Selected high-scoring TF–DORC pairs are labeled. **(B)** As in (A), for mCG–RNA FigR analysis. **(C)** Scatter plot of TF RNA expression–pseudotime Spearman correlation (x-axis) versus TF ATAC motif score–pseudotime Spearman correlation (y-axis) in EN L2/3 IT lineage cells. Each point represents one TF. Point size and opacity reflect $-\log_{10}$(BH-adjusted *p*-value) of the ATAC correlation. TFs with the strongest joint correlations are labeled. **(D)** As in (C), for TF mCG motif score-pseudotime Spearman correlation (y-axis). Point size and opacity reflect $-\log_{10}$(BH-adjusted *p*-value) of the mCG correlation. **(E)** ATAC-based TF–DORC regulatory network for the top activator and repressor TFs within the EN L2/3 IT lineage. TF node size indicates the number of regulated DORCs. TF node color reflects the ATAC motif score-pseudotime correlation direction. Node and edge opacity reflect their absolute correlation strength. Edge color indicates the regulation score direction, and width reflects the absolute regulation score. DORC nodes are shown in pink. **(F)** As in (E), for the mCG-based TF–DORC regulatory network.

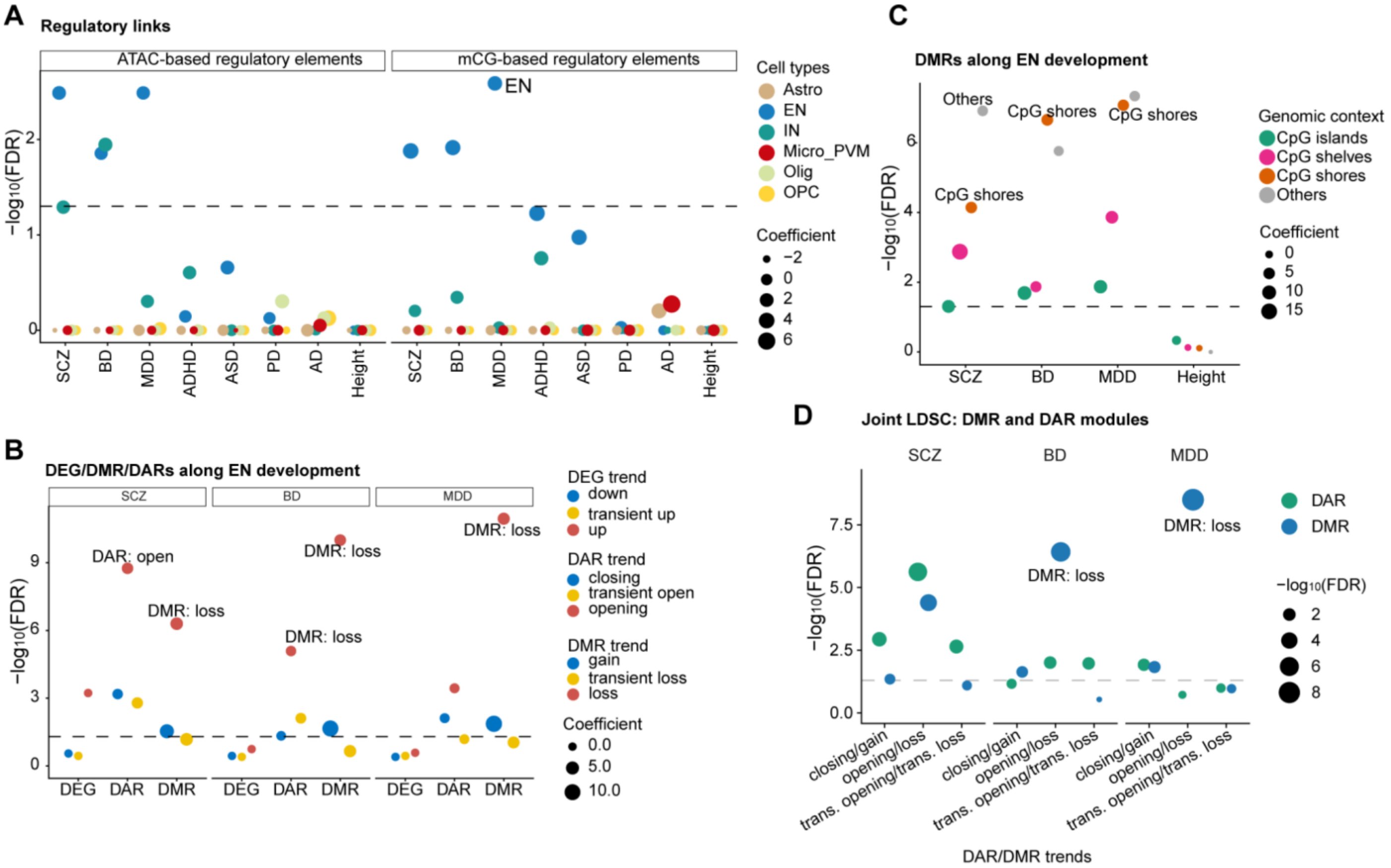


**Figure S9. Developmental epigenomic programs reveal complementary regulatory contributions to neuropsychiatric genetic risk, related to Figure 7. (A)** Stratified LD score regression (S-LDSC) analysis of neuropsychiatric and neurological disorder heritability enrichment across cell-type-specific regulatory elements defined by ATAC- and mCG-based regulatory links. **(B)** Significance of heritability enrichment of neuropsychiatric disorders across developmental DEGs, DARs, and DMRs. **(C)** Significance of heritability enrichment of neuropsychiatric disorders across DMRs stratified by CpG context, including CpG islands, shores, shelves, and other genomic regions. **(D)** Pairwise conditional S-LDSC analyses of corresponding developmental DAR and DMR modules jointly modeled alongside baseline annotations to assess their conditional contributions to trait heritability. Dashed horizontal lines indicate the significance threshold (*FDR* = 0.05).
